# CaMKII clusters neuronal LTCCs in biomolecular condensates to gate excitation-transcription coupling

**DOI:** 10.1101/2025.01.08.631979

**Authors:** Qian Yang, Xiaohan Wang, Lan Hu, Dorian Lawson-Qureshi, Roger J. Colbran

## Abstract

Neuronal depolarization activates L-type voltage-gated Ca²⁺ channels (LTCCs), increasing local Ca²⁺ concentrations to initiate excitation-transcription (E-T) coupling. We show that depolarization enhances clustering of Ca_V_1.2 and Ca_V_1.3 LTCCs in cultured hippocampal neurons, coinciding with increased nuclear CREB phosphorylation. LTCC clustering and LTCC-dependent CREB phosphorylation are selectively disrupted by 1,6-hexanediol, implicating biomolecular condensation. Activated CaMKIIα holoenzymes assemble complexes containing multiple Ca_V_1.2 and/or Ca_V_1.3 α1 subunits. Complex assembly is facilitated by co-expression of CaMKII-binding β2a subunits and Shank3 and selectively disrupted by 1,6-hexanediol. In HEK293 cells, pharmacological LTCC activation enhances clustering only when wild-type CaMKIIα is co-expressed. A CaMKII mutant that cannot bind LTCC N-terminal domains fails to support LTCC subunit complex formation *in vitro* and LTCC clustering in HEK293 cells. In neurons, the knockdown of CaMKII expression disrupts depolarization-induced (co-)clustering of Ca_V_1.2 and Ca_V_1.3. Together, these findings indicate that CaMKII-dependent clustering of plasma membrane LTCCs via biomolecular condensation is essential for initiating long-range signaling to activate gene expression following neuronal depolarization.

## Introduction

Voltage-gated calcium channels are protein complexes consisting of a pore-forming α1 subunit and auxiliary β, α2, and δ subunits. The auxiliary subunits modulate the intrinsic biophysical and regulatory properties of the α1 subunit, and influence cell surface targeting (Richards, Butcher and Dolphin, 2004; Voigt *et al*., 2016; Birnbaumer *et al*., 1998). The primary neuronal LTCC α1 subunits are Ca_V_1.2 and Ca_V_1.3, which exhibit highly overlapping expression patterns (Striessnig *et al*., 2014) and are often co-expressed in the same neuron. Although Ca_V_1.3 is activated preferentially with more modest membrane depolarization (Koschak *et al*., 2001; Xu and Lipscombe, 2001), Ca_V_1.2 and Ca_V_1.3 generally appear to regulate similar neuronal processes, such as excitation-transcription (E-T) coupling (Hetzenauer *et al*., 2006).

Learning and long-term memory require new gene transcription, which can be stimulated by many neurotransmitters. More specifically, LTCC-dependent E-T coupling links neuronal depolarization to the activation of several transcription factors that increase gene transcription (Ma *et al*., 2023; Bading, 2013; Yap and Greenberg, 2018), and can be initiated by locally elevated calcium concentrations near LTCCs without requiring global calcium increases in the cytosol or nucleus (Wheeler *et al*., 2012). It seems intuitive that the size and temporal dynamics of calcium nanodomains might be an important determinant of their biological function. Single LTCCs can create local calcium nanodomains (Tadross, Tsien and Yue, 2013; Nakamura, Reva and DiGregorio, 2018), but molecular mechanisms controlling calcium nanodomain dynamics are unclear. Previous studies revealed that LTCCs form clusters in the neuronal plasma membrane by poorly defined mechanisms, which facilitates the cooperative opening of some LTCC variants (Dixon *et al*., 2012; Moreno *et al*., 2016) and potentially modifies calcium nanodomain characteristics (Pfeiffer *et al*., 2020). Consequently, elucidating the molecular basis of LTCC clustering should provide new insights into the formation and function of calcium nanodomains.

Although LTCCs have an intrinsic ability to form clusters (Moreno *et al*., 2016; Dixon *et al*., 2012), clustering may also be modulated by additional proteins that interact with α1 or β auxiliary subunits (Weiss and Zamponi, 2017). For example, the postsynaptic scaffolding proteins Shank3 and densin modulate LTCC intracellular trafficking and/or calcium influx (Zhang *et al*., 2005; Jenkins *et al*., 2010; Wang *et al*., 2017a). Recent work showed that Shank3 promotes the clustering of neuronal Ca_V_1.3 α1 subunits under basal conditions (Yang *et al*., 2023). Prior studies have also shown that activated calcium/calmodulin-dependent protein kinase II (CaMKII), a highly abundant postsynaptic protein kinase, can interact directly with multiple intracellular domains of the Ca_V_1.2 or Ca_V_1.3 α1 subunits (Hudmon *et al*., 2005; Wang *et al*., 2017b; Simms *et al*., 2015), with β1 or β2a subunits (Grueter *et al*., 2008; Abiria and Colbran, 2010; Koval *et al*., 2010), and also with densin (Strack *et al*., 2000; Jiao *et al*., 2011; Walikonis *et al*., 2001; Jenkins *et al*., 2010; Wang *et al*., 2017a) and Shank3 (Perfitt *et al*., 2020; Cai *et al*., 2021). Some of these interactions have been shown to enhance CaMKII-dependent facilitation of calcium entry and/or E-T coupling. Pharmacological and molecular studies provided robust evidence that activated CaMKII is required for LTCC-dependent E-T coupling, at least in part by binding to RKR motifs in the N-terminal domain of Ca_V_1.3 and in a central domain of Shank3 (Wheeler *et al*., 2008; Ma *et al*., 2014; Wang *et al*., 2017b; Perfitt *et al*., 2020). However, the specific role of CaMKII in LTCC clustering remains unknown.

The ability of CaMKII holoenzymes to simultaneously interact with multiple proteins was initially recognized two decades ago (Robison *et al*., 2005). Some of these interactions have since been shown to confer a key role for CaMKII as a postsynaptic scaffolding protein, in addition to its conventional protein kinase function (Nicoll and Schulman, 2023; Bayer and Schulman, 2019). Indeed, both roles are crucial for the induction and maintenance of long-term potentiation at excitatory synapses (Rumian *et al*., 2024; Tullis *et al*., 2023; Chen *et al*., 2024).

Moreover, activated CaMKII has been shown to partition into phase-separated biomolecular condensates created *in vitro* by pre-mixing purified protein domains from one or more postsynaptic scaffolding proteins, including Shank3 (Hosokawa *et al*., 2021; Hayashi *et al*., 2021; Cai *et al*., 2021; Cai *et al*., 2023). Liquid-liquid phase separation and other forms of biomolecular condensation have emerged as potentially wide-spread mechanisms for membrane-less compartmentalization of molecular processes in cells (Gao *et al*., 2022; Wang *et al*., 2021; Mehta and Zhang, 2022), including at neuronal synapses (Guzikowski and Kavalali, 2024; Zeng *et al*., 2016; Zeng *et al*., 2018; Zeng *et al*., 2019; Milovanovic *et al*., 2018).

Here, we show that depolarization enhances the clustering of Ca_V_1.3 and Ca_V_1.2 LTCCs in cultured hippocampal neurons, and that clustering is disrupted by 1,6-hexanediol, which is often used to disrupt biomolecular condensates (Zheng *et al*., 2025; Kato, Zhou and McKnight, 2022). Moreover, 1,6-hexanediol also interferes with neuronal E-T coupling. CaMKII facilitates the homo-or hetero-meric clustering of Ca_V_1.3 and Ca_V_1.2 LTCC complexes in an activation-dependent manner *in vitro* and on the membrane of HEK293 cells. This LTCC clustering requires direct interaction of CaMKII with Ca_V_1.3 or Ca_V_1.2 and is facilitated by β2a auxiliary subunits and Shank3. Our data indicate that depolarization-dependent Ca_V_1.3 and Ca_V_1.2 LTCC clustering, as well as Ca_V_1.3/Ca_V_1.2 co-clustering, is disrupted by suppressing CaMKII expression in neurons. Collectively, these findings indicate that the initiation of E-T coupling following neuronal membrane depolarization involves the activated CaMKII-dependent assembly of multimeric LTCC complexes in biomolecular condensates.

## Method

### DNA constructs

To generate FLAG-Ca_V_1.3, the DNA sequence encoding HA in the pCGN vector was first deleted to yield pCGN0 by mutagenesis. A double-stranded DNA oligo linker that encodes FLAG (Forward: CT AGC **GAC TAC AAA GAC GAT GAC GAC AAG** TCT AGA GGC G; Reverse: GA TCC GCC TCT AGA **CTT GTC GTC ATC GTC TTT GTA GTC** G; sequences encoding the FLAG epitope are in bold) was then inserted into pCGN0 to yield pCGNF. The cDNA sequence of Ca_V_1.3 was then inserted into pCGNF between the XbaI and BamHI sites using SLIC cloning. To make pcDNA-msCaMKII-ΔAD (association domain truncated), the DNA sequence encoding E341 to H478 was deleted from the pcDNA-msCaMKII construct. All deletions were done following the one-step mutagenesis protocol described by Liu et al (Liu and Naismith, 2008). The original sources of all DNA constructs are provided in the Key Resources Table. All constructs were confirmed by DNA sequencing.

### HEK cell culture and transfection

HEK293 and HEK293T cells (purchased from ATCC) were cultured and transfected as previously described (Yang *et al*., 2023). Briefly, cells were grown in high glucose DMEM at 37°C, 5% CO2, and passaged every 3-4 days. Cells (≤ 20 passages) were transfected using Lipofectamine 2000 after reaching approximately 70% confluence. For Figure S2, the ratio of α1: α2δ: β3: CaMKIIα was set at was set at 2:1:1:1. The ratio of iHA-Ca_V_1.3: FLAG-Ca_V_1.3 was set at 1:1. For the rest of co-immunoprecipitation experiments, tagged Ca_V_1.3 or/and Ca_V_1.2 α1 subunits were always co-expressed with α2δ and either β3 or β2a with a FLAG tag (FLAG-β3 or -β2a) auxiliary subunits, together with pcDNA vector (empty or encoding CaMKIIα). The ratio of α1: α2δ: FLAG-β:pcDNA was set at 3:1:1:1.2. Co-transfections with N-terminal intracellular HA-tagged Ca_V_1.3 (iHA-Ca_V_1.3): and either mCherry-Ca_V_1.3 or GFP-Ca_V_1.2 used a DNA ratio of 1:1.5. Co-transfections involving GFP-Shank3 or CaMKII, or a combination of GFP-Shank3 and CaMKIIα, with LTCC subunits were performed using a DNA ratio of 3:1:1:1:0.5 (α1:α2δ:FLAG-β2a:CaMKII:Shank3).

When GFP-Shank3 or CaMKIIα was expressed individually, empty pCDNA or GFP vector was incorporated to maintain the same overall DNA quantity as co-transfection with GFP-Shank3 and CaMKIIα. GFP-Shank3 and mApple-Shank3 or CaMKII, were co-expressed using a DNA ratio of 1:2 (GFP-Shank3:mApple-Shank3/CaMKII). For co-immunoprecipitation experiments, HEK293T cells were transfected with a maximum of 10 μg DNA per 10-cm dish and harvested ∼48 hours later. For immunostaining experiments, HEK293 cells were transfected with a maximum of 2 μg DNA per well of 6-well plates, and re-plated onto coverslips in 24-well plates ∼24 hours later. Coverslips were used for experiments after another 24-36 hours and then fixed.

### Co-immunoprecipitation

Transfected HEK 293T cells were lysed and used for co-immunoprecipitation using HA antibody or GFP antibody and magnetic Dynabeads Protein A beads as previously described (Yang *et al*., 2023). Where indicated, lysates were supplemented with 2 mM CaCl_2_ and/or 1 µM calmodulin (Ca^2+^/calmodulin), either alone or with 50 µM calmidazolium (Figure S4), prior to incubation.

Some experiments included the indicated concentrations of 1,6-hexanediol, added to the lysates simultaneously with (or without) Ca^2+^/calmodulin. Only in Figure S2, lysates were supplemented with or without 2 mM CaCl_2_, 1 µM calmodulin, 2 mM MgCl_2_, and 1 mM ATP.

### Western blotting

Co-immunoprecipitation samples were separated on either 7.5% (Figures 2A, 2E, 2G, S3) or 10% (Figures 2C, 3, 4, S2, S4-6) SDS-PAGE gels, followed by transfer to nitrocellulose membrane as previously described (Yang *et al*., 2023). Membranes were stained with Ponceau-S to confirm protein transfer and loading, and digitally scanned before being incubated in blocking buffer (5% nonfat milk in Tris-buffered saline with 0.1% (v/v) Tween-20 (TBST)) for one hour at room temperature. Primary antibodies (rabbit anti-HA, mouse anti-CaMKII, and rabbit anti-Shank3 were diluted 1:8000; all other primary antibodies were diluted 1:4000) in blocking buffer, and incubated with membranes overnight at 4°C. Note that GFP eluted from the bottom of 7.5% gels and so GFP blots are not shown for these experiments. After washing three times (10 minutes per wash) with TBST, membranes were incubated with IR dye-conjugated secondary antibodies (donkey anti-rabbit 680LT diluted 1:8000 and donkey anti-mouse 800CW diluted 1:4000) or with HRP-conjugated secondary antibodies (rat anti-mouse) in blocking buffer for one hour at room temperature. After two further washes, membranes were scanned using an Odyssey system and images were quantified by Odyssey Image Studio Software. Identically sized rectangles were drawn to select a specific protein band for quantification across all the lanes on a gel and the background was subtracted using the median background method.

Specifically, the area used to compute background is taken from a three-pixel border around top and bottom sides of each rectangle. Membranes incubated with HRP-conjugated secondary antibodies were incubated with the Western Lightening Plus-ECL, enhanced chemiluminescent substrate (PerkinElmer) and visualized using Premium X-ray Film (Phenix Research Products) exposed in the linear response range.

### HEK cell stimulation and immunocytochemistry

HEK293 cells expressing iHA-Ca_V_1.3 LTCCs with pcDNA vector or CaMKII (WT or V102E) were pre-incubated in Ca^2+^-free HEPES buffer (150 mM NaCl, 5 mM KCl, 2 mM MgCl_2_, 10 mM HEPES pH 7.4, 10 mM Glucose) for 10 min, and then transferred to HEPES buffer supplemented with 10 μM BayK-8644 (BayK) and/or 2.5 mM CaCl_2_ for 5-10 min. DMSO (0.02% v/v) was added to incubations lacking BayK as a vehicle control. Cells were then fixed using ice-cold 4% paraformaldehyde containing 4% sucrose in 0.1 M Phosphate Buffer (pH 7.4) (4% PFA) for 10 min at room temperature, washed three times (5 min per wash) with PBS, and then permeabilized with PBS containing 0.2% Triton X-100 for 8 min. Subsequently, cells were incubated with blocking solution (1X PBS, 0.1% Triton X-100 (v/v), 2.5% BSA (w/v), 5% Normal Donkey Serum (w/v), 1% glycerol (v/v)) at room temperature for 1 h, then incubated with primary antibodies (rabbit anti-HA diluted 1:1000 and mouse anti-CaMKII diluted 1:2500) in blocking buffer overnight at 4°C. After three washes with PBS containing 0.2% Triton X-100 (10 min each), cells were incubated for one hour at room temperature with secondary antibodies (donkey anti-rabbit Alexa Fluor 647 and donkey anti-mouse Alexa Fluor 488), each diluted 1:2000 in blocking buffer. After three additional washes with PBS, coverslips were mounted on slides using Prolong Gold Antifade Mountant and then stored at 4°C for subsequent imaging.

### Total internal reflection fluorescence (TIRF) microscopy and quantification

A Nikon Multi Excitation TIRF microscope with 488 nm and 640 nm solid-state lasers was used as described (Yang *et al*., 2023). The automatic H-TIRF module was used for the 488 laser and the manual TIRF module was used for the 640 laser. Calibration of the incident angle and focus for achieving TIRF was conducted individually for each module. Images were then acquired using NIS-Elements. Consistency was maintained by using the same exposure time of 80–150 ms for both channels, alongside laser power at 5-8% for 488 nm laser and 10–12% of 640 nm laser. These imaging parameters remained uniform across all cover slips within the same biological replicate. Images were processed in Fiji software (ImageJ, NIH) to quantify cluster density and intensity of surface localized iHA-Ca_V_1.3, as well as to assess colocalization between iHA-Ca_V_1.3 and CaMKII. Specifically, cells positive with both iHA and CaMKII signal were selected to be quantified, using freehand selections. These ROIs were then consistently applied to the iHA-Ca_V_1.3 channel after background subtraction. A threshold for the HA signal was established using the mean intensity of HA signal plus 1.5 times the standard deviation. The “Analyze Particles” function was used to calculate the intensity, area, and count of iHA-Ca_V_1.3 clusters surpassing the threshold within each ROI. Cluster density was derived by dividing the cluster count by the area of the respective ROI. For colocalization analysis, both CaMKII and HA channels underwent automated thresholding before evaluating the intensity correlation quotient (ICQ). The metric quantifies colocalization on a scale from −0.5 to +0.5, delineating the extent from complete segregation to perfect overlap, as previously described (Li *et al*., 2004; Perfitt *et al*., 2020; Yang *et al*., 2023).

### Primary hippocampal neuron cultures and depolarization

Dissociated hippocampal neurons were prepared from E18 Sprague Dawley rat embryos, as previously described (Shanks *et al*., 2010) resulting in cultures containing an approximate 50:50 mix of neurons from male and female embryos. At 14 days *in vitro* (DIV), neurons on coverslips in 12 well plates were co-transfected using Lipofectamine 2000 (Thermo Fisher Scientific) and 1 µg DNA, in accordance with the manufacturer’s directions, to express Ca_V_1.3/Ca_V_1.2 α1 subunit with a N-terminal intracellular HA tag (iHA-Ca_V_1.3) or an extracellular HA tag (sHA-Ca_V_1.3/sHA-Ca_V_1.2), FLAG-β2a or -β3, and GFP-nonsense shRNA (nssh) or GFP-CaMKIIα-shRNA and GFP-CaMKIIβ-shRNA (GFP-CaMKII-shRNA) at an α1: β: GFP ratio of 3:1:1.2. After a further 7 days, neurons were pre-incubated in 5K Tyrode’s solution (150 mM NaCl, 5 mM KCl, 2 mM CaCl_2_, 2 mM MgCl_2_, 10 mM glucose and 10 mM HEPES pH 7.5 (∼313 mOsm)) with inhibitors (1 μM TTX, 10 μM APV, and 50 μM CNQX) for 2 hours at 37°C and 5% CO_2_ to inhibit the intrinsic neuronal activity. Neurons were then incubated with either 5K or 40K Tyrode’s solution (adjusted to 40 mM KCl and 115 mM NaCl, with inhibitors present) with or without the addition of 1,6-hexanediol for 90 seconds and then quickly fixed in ice-cold 4% PFA for 10 minutes at room temperature, followed by three PBS washes. For Forskolin treatment, neurons at DIV 21 were pre-incubated with inhibitors (1 μM TTX, 10 μM APV, 50 μM CNQX, and 10 μM Nimodipine) for 30-60 min at 37°C and 5% CO2 to inhibit intrinsic neuronal activity. Half of the culture medium was then replaced with fresh medium containing the same inhibitors and one of the following treatments: vehicle control, 150 μM Forskolin (final concentration 75 μM), or Forskolin combined with 2% 1,6-or 2,5-Hexanediol (final concentration 1%). After a further 5 min incubation, neurons were rapidly fixed in ice-cold 4% PFA for 10 min at room temperature, followed by three washes with PBS. For surface sHA-Ca_V_1 staining, neurons were blocked in non-permeabilizing blocking buffer A (1X PBS, 5% (w/v) bovine serum albumin (Sigma)) for 1 hour. Primary anti-HA antibody (rabbit or mouse, diluted 1:200 in blocking buffer A) was applied overnight at 4°C. After three PBS washes (10 min per wash), neurons were incubated with secondary antibodies (donkey anti-rabbit or anti-mouse Alexa Fluor 647, diluted 1:200 in blocking buffer A) for 1 hour at room temperature. Following three additional washes with PBS, neurons were permeabilized by PBS containing 0.2% Triton X-100 for 8 min, then blocked using blocking buffer B (1X PBS, 0.1% Triton X-100 (v/v), 2.5% BSA (w/v), 5% Normal Donkey Serum (w/v), 1% glycerol (v/v)) for 1 hour. For the following staining for rabbit anti-pCREB (dilution: 1:1000) or mouse anti-CaMKII (dilution: 1:2000), the process was the same as the permeabilized immunostaining in HEK293 cells.

### Neuronal imaging and quantification

Neuronal imaging was conducted using Zeiss LSM 880 microscope with a 63x/1.4 Plan-APOCHROMAT oil lens as previously described (Yang *et al*., 2023). Transfected neurons were identified through GFP signals under a binocular lens. Conventional confocal mode was used in Figures 1B-C, 8, and S7-S8. For whole cell imaging (Figures 1C and S7A), Z-stack images were captured with a step size of 0.3 μm and a range of 1.8-2.7 μm. Z-stack images were then processed in Fiji software to create a maximum intensity projection for subsequent analysis. To collect pCREB images, the DAPI channel was used to adjust the focus of nucleus. In Figures 9-12, the Airyscan mode was used to maximize sensitivity and resolution for the collection of surface-localized sHA-Ca_V_1.2/sHA-Ca_V_1.3 and endogenous CaMKII/Ca_V_1.2 images, collecting single focal plane images (73.51 × 73.51 μm) (Yang *et al*., 2023).

**Figure 1.**
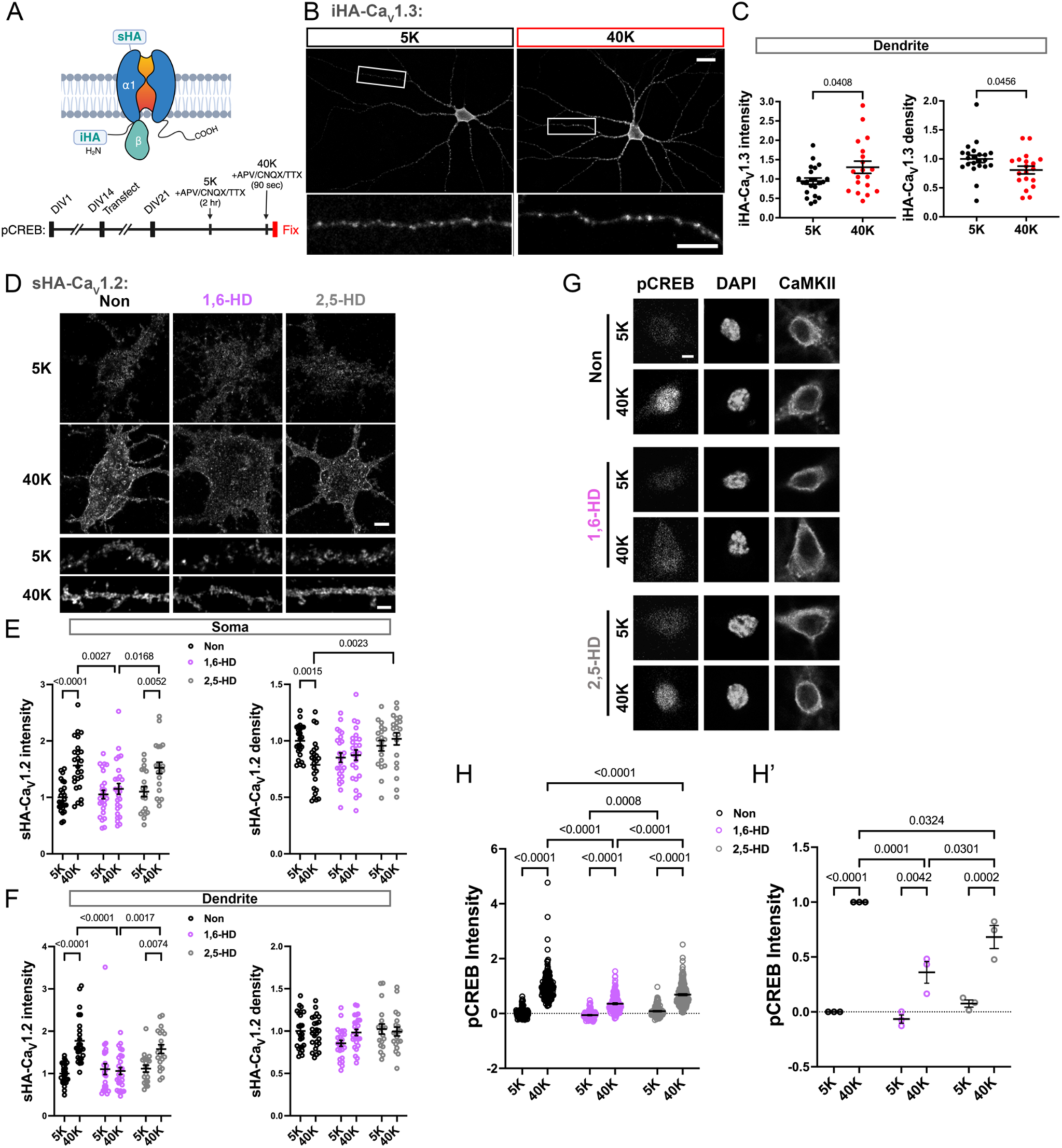
Activity-dependent regulation of dendritic LTCC clustering is selectively disrupted by 1,6-hexanediol in hippocampal neurons. A) Schematic of neuronal transfection and experimental protocols. Rat primary hippocampal neurons were co-transfected with HA-tagged LTCC α1 subunits (intracellular tag: iHA-Ca_V_1.3 or extracellular tag: sHA-Ca_V_1.2 or sHA-Ca_V_1.3) and FLAG-β2a at 14 days in vitro (DIV 14). At DIV 21, neurons were pre-incubated in 5K Tyrode’s solution for 2 h and then stimulated for 90 s with either 5K or 40K Tyrode’s solution. Neurons were immediately fixed, permeabilized and stained as described below. Details provided in Methods. B) Representative images of iHA-Ca_V_1.3 staining in whole neurons (above) and dendrites (bottom) after 5K or 40K treatment. Scale bars: 20 μm for whole cells, 10 μm for dendrites. C) Quantification of relative iHA-Ca_V_1.3 cluster density and cluster intensity, respectively, in dendrites of n = 22 (5K) or 19 (40K) neurons from three independent cultures. Statistical analyses: unpaired t-tests. D) Representative images of sHA-Ca_V_1.2 staining in soma and dendrites of neurons (also expressing soluble GFP; not shown) incubated for 90 s with 40K or 5K with or without 1% 1,6- or 2,5-hexanediol (HD) and then fixed. HA immunostaining was prior to permeabilization. Images collected using Airyscan super-resolution confocal microscopy. Scale bar, 5 μm. E-F) Quantification of sHA-Ca_V_1.2 cluster intensity and density in soma (B) and dendrites (C) of n = 25 (5K), 25 (40K), 25 (5K with 1,6-HD), 26 (40K with 1,6-HD), 19 (5K with 2,5-HD), and 20 (40K with 2,5-HD) GFP expressing neurons. Data were quantified from 3 independent cultures/transfections and normalized to the mean values in neurons following the 5K incubation without HD application within each replicate. The 2,5-HD treatment was included in only two of the replicates included in panels E-F. Statistical analyses: two-way ANOVA followed by Šídák’s post hoc tests. G) Representative images of pCREB, DAPI, and CaMKII immunostaining in hippocampal neurons (DIV 21) incubated for 90 s with 40K/5K with or without 1% 1,6- or 2,5-hexanediol (HD) and then fixed. Images were collected using confocal microscopy. Scale bar, 5 μm. H) Quantification of pCREB signal after 5K or 40K treatment in neurons with or without 1,6-HD or 2,5-HD application (Non: n = 210 for 5K and 173 for 40K; 1,6-HD: n = 132 for 5K and 191 for 40K; 2,5-HD: n = 182 for 5K and 211 for 40K). H’) Re-analysis of data in panel H by the 3 replicates. Data were normalized to the mean for neurons following the 40K incubation without HD within each replicate. Statistical analyses: two-way ANOVA with Tukey’s post hoc tests.

All image quantifications were performed in Fiji software. For endogenous CaMKII analysis, CaMKII signal in the soma was designated as regions of interest (ROIs) in both GFP-positive or - negative neurons. Background was subtracted, and the mean intensities of CaMKII signal in the ROIs were measured. For pCREB analysis, ROIs were assigned based on DAPI signals, and pCREB intensities in these ROIs from GFP-positive or -negative neurons were measured. For LTCC cluster analysis, ROIs were selected based on the GFP channel. Somatic ROIs were selected similarly to that in HEK293 cells, while dendritic ROIs were selected followed rules outlined previously (Yang *et al*., 2023). Background was then subtracted, sHA signals were thresholded, and the Analyze Particles function was used, as in HEK293 cell analyses. For colocalization between sHA-Ca_V_1.3 and endogenous Ca_V_1.2, soma and dendritic ROIs were identified based on GFP signals. Following background subtraction, the average intensities of sHA-Ca_V_1.3 (x̄_1.3_) and Ca_V_1.2 (x̄_1.2_) in ROIs were measured. We then defined new ROIs based on sHA staining and separately measured the intensity of sHA-Ca_V_1.3 and Ca_V_1.2 in these ROIs. After normalizing sHA signal to x̄_1.3_ and normalizing Ca_V_1.2 signal to x̄_1.2_, the ratio of normalized Ca_V_1.2 to normalized sHA intensity in somatic and dendritic ROIs were used as a measure of endogenous Ca_V_1.2 colocalization with sHA-Ca_V_1.3. The experimenter was blind to the experimental conditions when analyzing (i/s)HA-Ca_V_1.3/ sHA-Ca_V_1.2 clusters.

## Statistical analysis

All data are shown as mean ± SEM, and sample sizes shown as n refer to the number of cells or independent biological replicates (experiments) as indicated in figure legends. All statistical tests were performed in GraphPad Prism 8 software (GraphPad). Differences were considered significant if p ≤ 0.05. For comparisons between two groups, unpaired Student’s t-test (two-tailed) was used. For comparisons between three or more groups with two independent variables, two-way ANOVA followed by the post hoc tests recommended by Prism was used (defined in each relevant figure legend).

## Results

### Neuronal depolarization increases Ca_V_1.3 LTCC clustering

We recently reported that Shank3 clusters Ca_V_1.3, but not Ca_V_1.2, LTCCs *in vitro*, in HEK293 cells and in cultured hippocampal neurons under basal conditions, and that clustering can be disrupted by Ca^2+^ *in vitro* and in HEK293 cells (Yang et al, 2023). To test the hypothesis that Ca^2+^ influx disrupts neuronal Ca_V_1.3 LTCC clustering, cultured hippocampal neurons expressing N-terminal intracellular HA-tagged Ca_V_1.3 (iHA-Ca_V_1.3), α2δ, and FLAG-β2a (DIV21; 7 days post-transfection) were briefly (90 s) depolarized using a well-established 40 mM KCl (40K) protocol in the presence of APV, CNQX and TTX to block NMDA receptors, AMPA receptors, and voltage-dependent sodium channels, respectively (Figure 1A). This stimulation paradigm triggers E-T coupling in DIV14 hippocampal neurons, as measured by phosphorylation of the CREB transcription factor and by expression of the c-fos immediate early gene (Wheeler *et al*., 2008; Wang *et al*., 2017b; Perfitt *et al*., 2020). In parallel, the 40K treatment significantly increased the intensity and significantly decreased the density of dendritic iHA-Ca_V_1.3 clusters relative to the control 5K basal treatment (Figure 1B-C). These results indicate that Ca^2+^ influx following neuronal depolarization to initiate E-T coupling further enhances Ca_V_1.3 LTCC clustering, inconsistent with our initial hypothesis.

### Involvement of biomolecular condensation in depolarization-induced LTCC clustering

Biomolecular condensation is emerging as a fundamental mechanism for the membrane-less compartmentalization of intracellular proteins to efficiently perform diverse biological functions (Gao *et al*., 2022; Mehta and Zhang, 2022; Wang *et al*., 2021). Indeed, recent data indicate that biomolecular condensates play key roles in the organization of both pre- and post-synaptic proteins at excitatory synapses (Guzikowski and Kavalali, 2024; Zeng *et al*., 2016; Zeng *et al*., 2018; Zeng *et al*., 2019; Milovanovic *et al*., 2018) and were recently shown to modulate the distribution of presynaptic voltage-gated Ca^2+^ channels (Jin *et al*., 2025). Therefore, we extended our analysis to Ca_V_1.2 LTCCs and tested for the involvement of biomolecular condensates in depolarization-induced LTCC clustering by comparing the effects of 1,6-hexanediol or 2,5-hexanediol during neuronal depolarization. We used 2,5-hexanediol as a control because biomolecular condensates are often more sensitive to disruption by 1,6-hexanediol compared to 2,5-hexanediol due to differential spacing of the two hydroxyl moieties on the hexane backbone (Kato, Zhou and McKnight, 2022).

Cultured hippocampal neurons expressing extracellular HA-tagged Ca_V_1.2 (sHA-Ca_V_1.2), α2δ, and FLAG-β2a (DIV21; 7 days post-transfection) were incubated under basal conditions (see above) and then depolarized for 90 s with 40K Tyrodes solution in the absence or presence of 1,6- or 2,5-hexanediol (1% v/v each). Note that hexanediols were present only during the 90 s depolarization. After immediate fixation and staining, we quantified the intensity and density of surface-localized sHA-Ca_V_1.2 clusters. Under basal (unstimulated) conditions, the intensities and densities of sHA-Ca_V_1.2 staining in the soma and dendrites were unaffected by either hexanediol (Figure 1D-F). However, depolarization significantly increased sHA-Ca_V_1.2 cluster intensities in both the somatic and dendritic compartments, as seen for iHA-Ca_V_1.3 (see above). Morever, 1,6-hexanediol disrupted the depolarization-induced increases in staining intensity, while 2,5-hexanediol had no effect (Figure 1D-F and S1). These results support the hypothesis that depolarization-induced neuronal LTCC clustering involves biomolecular condensation.

### 1,6-Hexanediol disrupts E-T coupling

As an initial test of the relationship between 1,6-hexanediol-sensitive LTCC clustering and E-T coupling in cultured hippocampal neurons, we investigated the sensitivity of depolarization-induced CREB Ser133 phosphorylation to 1,6- and 2,5-hexanediol (1% v/v each) (Figure 1G-H). Neither hexanediol has a significant effect on low levels of CREB phosphorylation under basal conditions. However, inclusion of 1,6-hexanediol during the 40K depolarization significantly reduced CREB phosphorylation by approximately 64% compared to the control in the absence of hexanediol. Notably, 2,5-hexanediol had a significantly weaker effect, reducing CREB phosphorylation by only ∼31% under the same conditions (Figure 1H–H′). As an additional control, we found that neither 1,6- nor 2,5-hexanediol had a significant effect on cyclic AMP-stimulated neuronal CREB phosphorylation (Figure S2). The significantly stronger effect of 1,6-hexanediol on depolarization-induced CREB phosphorylation supports the hypothesis that biomolecular condensation is involved in the E-T coupling signaling pathway that leads to CREB phosphorylation.

### Activated CaMKII assembles complexes containing multiple Ca_V_1.3 LTCCs

Binding of activated CaMKII catalytic domains to the N-terminal domain of Ca_V_1.3 LTCC α1 subunits is required for E-T coupling (Wheeler *et al*., 2012; Ma *et al*., 2014; Wang *et al*., 2017b). Moreover, we found that this interaction enhances the immunoprecipitation of iHA-Ca_V_1.3 from transfected HEK293T cell lysates by an HA antibody (Wang *et al*., 2017b). As an initial test of the hypothesis that the increased iHA-Ca_V_1.3 immunoprecipitation efficiency may be due to simultaneous binding of multiple iHA-tagged α1 subunits to subunits within activated CaMKII holoenzymes, we co-expressed iHA-Ca_V_1.3 with β3 and α2δ subunits and wild type (WT) or truncated/monomeric CaMKIIα in HEK293T cells and immunoprecipitated iHA-CaV1.3 from lysates. Compared to immunoprecipitation under basal (EDTA) conditions, the addition of Ca^2+^/calmodulin/Mg^2+^-ATP to the lysates resulted in a 4-5-fold (n=2) increase in the amount of iHA-CaV1.3 that was immunoprecipitated when lysates contained WT CaMKIIα. Deletion of residues 69-93 in the iHA-CaV1.3 N-terminal domain (Δ69-93), preventing the direct interaction with CaMKIIα, blocked the Ca^2+^/calmodulin/Mg^2+^-ATP-induced increase of iHA-CaV1.3 immunoprecipitation, consistent with our prior report (Wang *et al*., 2017b). We now show that the increased immunoprecipitation of iHA-Ca_V_1.3 following Ca^2+^/calmodulin/Mg^2+^-ATP addition also was not observed in the presence of a monomeric CaMKIIα (created by truncation of the CaMKII association domain at residue 340 (ΔAD)). However, the addition of Ca^2+^/calmodulin/Mg^2+^-ATP increased the amount of co-precipitated ΔAD-CaMKIIα and WT CaMKIIα, confirming that both the monomeric kinase and WT kinases bind similarly to iHA-CaV1.3 (Figure S3A). These data suggest that the increased immunoprecipitation of WT iHA-Ca_V_1.3 by the HA antibody following Ca^2+^/calmodulin/Mg^2+^-ATP is due to simultaneous interaction of the N-terminal domains of multiple iHA-Ca_V_1.3 channels with different subunits within activated CaMKII holoenzymes.

To more directly test the hypothesis that activated CaMKII holoenzymes are capable of binding simultaneously to multiple Ca_V_1.3 α1 subunits, similar experiments were conducted using lysates of HEK293T cells co-expressing FLAG- and iHA-tagged Ca_V_1.3 α1 subunits (WT or mutated) and β3 and α2δ subunits, with or without CaMKIIα. Addition of Ca^2+^/calmodulin/Mg^2+^-ATP to lysates increased the coimmunoprecipitation of both WT CaMKIIα and FLAG-Ca_V_1.3 with iHA-Ca_V_1.3 (n=2), but these increases were largely prevented by mutations of iHA-Ca_V_1.3 (Δ69-93) (n=2) or CaMKIIα (V102E) (n=1) that disrupt their interaction (Figure S3B). These data further support the hypothesis that activated CaMKII holoenzymes can assemble complexes containing multiple Ca_V_1.3 α1 subunits.

Prior studies found that activated CaMKIIα also binds to LTCC β1 and β2a subunits, but not to β3 or β4 subunits (Grueter *et al*., 2008; Abiria and Colbran, 2010). Therefore, to test the hypothesis that β subunits also play a role in CaMKII-dependent assembly of multi-Ca_V_1.3 complexes, we compared HA-immunoprecipitions from lysates of HEK293T cells co-expressing iHA- and mCherry-Ca_V_1.3 α1 subunits with FLAG-tagged β2a or β3 subunit and the α2δ subunit, with or without CaMKIIα. Under basal lysate conditions (with EDTA present), both FLAG-tagged β subunits co-immunoprecipitated with the HA antibody, as expected, but negligible amounts of mCherry-Ca_V_1.3 α1 or CaMKIIα were detected. Initial studies found that the addition of Ca^2+^/calmodulin/Mg^2+^-ATP to lysates containing CaMKIIα resulted in substantial FLAG-β subunit phosphorylation (Abiria and Colbran, 2010), potentially confounding data interpretation.

Therefore, we activated CaMKIIα by adding Ca^2+^/calmodulin alone to the cell lysates, which significantly increased mCherry-Ca_V_1.3 co-immunoprecipitation with the HA-antibody, but only in the presence of co-expressed CaMKIIα (Figure 2A-B), similar to initial observations using FLAG-Ca_V_1.3 with untagged β3 subunit (Figure S3B). Notably, significant amounts of CaMKIIα co-immunoprecipitated with iHA-Ca_V_1.3 only following the addition of Ca^2+^/calmodulin.

**Figure 2.**
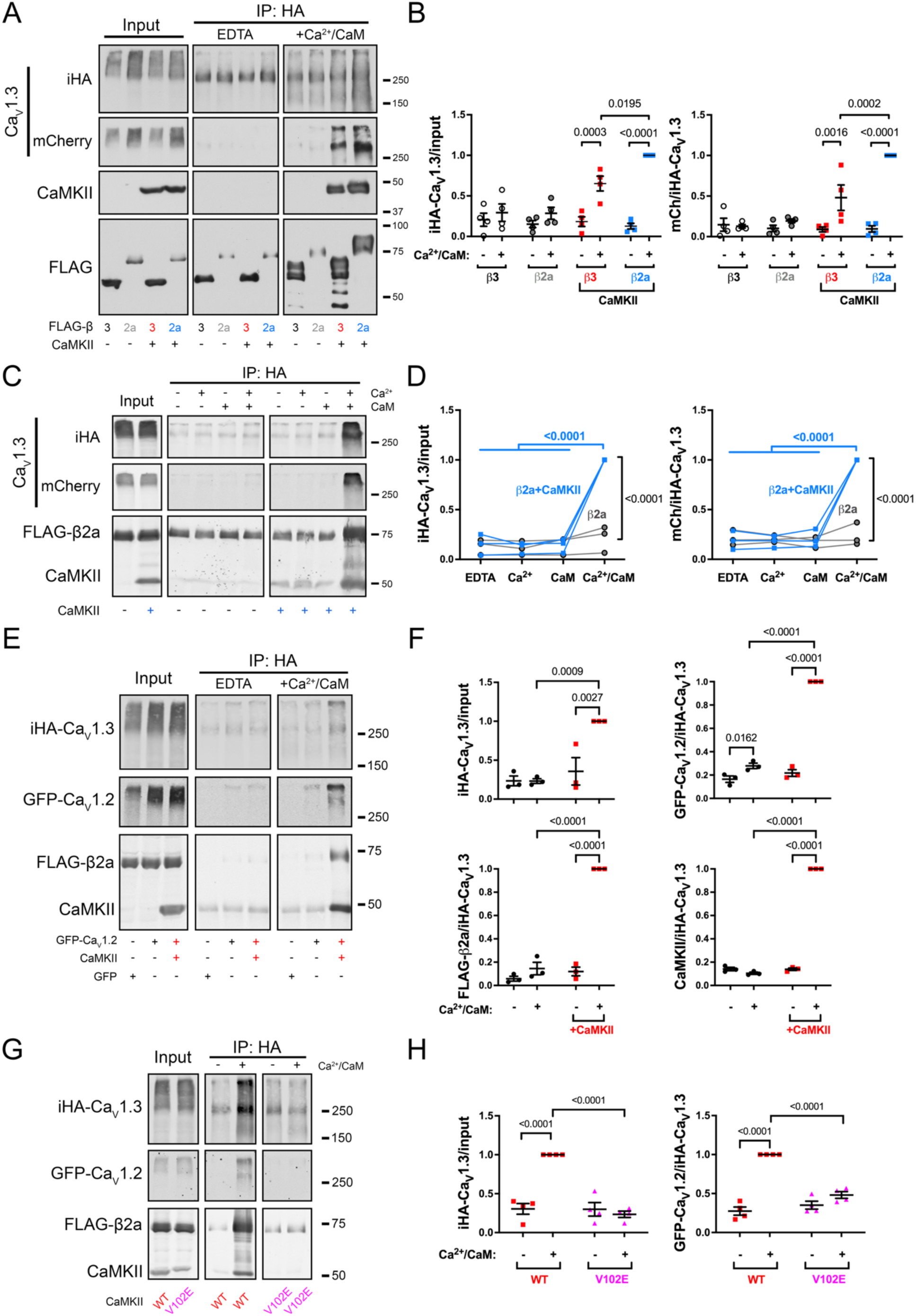
CaMKIIα- and Ca^2+^/calmodulin-dependent assembly of LTCC complexes in HEK293T cell lysates. A) Representative immunoblots of the input and anti-HA immunoprecipitations (IPs) from soluble fractions of HEK293T cells co-expressing iHA-Ca_V_1.3, mCherry-Ca_V_1.3, α2δ, FLAG-β3 or -β2a subunits, with or without CaMKIIα. Duplicate aliquots of the same lysates were immunoprecipitated with no further additions (EDTA) or following addition of Ca^2+^/calmodulin (Ca^2+^/CaM). B) Quantification of iHA-Ca_V_1.3 and mCherry-Ca_V_1.3 signals in HA-immune complexes from four independent transfection replicates. For each replicate, iHA-Ca_V_1.3 signals in the IP lanes were normalized to the corresponding input (iHA-Ca_V_1.3/input); mCherry-Ca_V_1.3 signals were normalized to iHA-Ca_V_1.3 in the corresponding lane (mCh/iHA-Ca_V_1.3). Ratios within each replicate were then normalized to the “Ca^2+^/calmodulin/β2a + CaMKII” condition to allow pooling of the data from all replicates. Statistical analyses: two-way ANOVA with Šídák’s post hoc tests. C) Representative immunoblots of the input and anti-HA IPs from soluble fractions of HEK293T cells co-expressing iHA-Ca_V_1.3, mCherry-Ca_V_1.3, α2δ, and FLAG-β2a subunits, with or without CaMKII. Aliquots of the same lysates were supplemented with the addition of nothing, Ca^2+^ alone, calmodulin alone, or both Ca^2+^ and calmodulin. D) Quantification of iHA-Ca_V_1.3 and mCherry-Ca_V_1.3 signals in HA-immune complexes from three independent transfection replicates. Immunoblot signals were normalized relative to the Ca^2+^/calmodulin + CaMKII condition within each replicate as described for panel B. Statistical analyses: two-way ANOVA with Šídák’s post hoc tests (for comparisons without and with CaMKII co-expression) or by Tukey’s post hoc tests (for comparisons between four conditions). E) Representative immunoblots of the input and anti-HA IPs from soluble fractions of HEK293T cells co-expressing iHA-Ca_V_1.3, GFP-Ca_V_1.2, α2δ, and FLAG-β2a subunits, with or without CaMKIIα, and without or with Ca^2+^/calmodulin addition. F) Quantification of iHA-Ca_V_1.3, GFP-Ca_V_1.2, FLAG-β2a, and CaMKIIα signals in HA-immune complexes from three independent transfection replicates. Immunoblot signals were normalized relative to the Ca^2+^/calmodulin condition with CaMKII co-expression as described for Panel B. G) Representative immunoblots of the input and anti-HA IPs from soluble fractions of HEK293T cells co-expressing iHA-Ca_V_1.3, GFP-Ca_V_1.2, α2δ, FLAG-β2a subunits, and CaMKIIα (WT or V102E mutant) without or with Ca^2+^/calmodulin addition. H) Quantification of iHA-Ca_V_1.3 and GFP-Ca_V_1.2 signals in HA-immune complexes from four independent transfection replicates. Immunoblot signals were normalized relative to the Ca^2+^/calmodulin condition with CaMKII-WT co-expression as described for Panel B. Statistical analyses: two-way ANOVA with Šídák’s post hoc tests. Images of immunoblots for each protein in panels A, C, E and G were from the same transfection replicate gel/blot and were collected using identical detection sensitivity.

Ca^2+^/calmodulin-enhanced co-immunoprecipitation of mCherry-Ca_V_1.3 and CaMKIIα with iHA-Ca_V_1.3 was substantially larger in the presence of FLAG-β2a relative to the presence of FLAG-β3, even though the cells expressed higher levels of FLAG-β3 than FLAG-β2a, as measured by the anti-FLAG immunoblot (Figure 2A-B). Taken together, these data confirm that CaMKIIα activation promotes CaMKIIα association with Ca_V_1.3 to assemble complexes containing multiple Ca_V_1.3 α1 subunits, and further show that CaMKIIα-dependent assembly of these complexes is facilitated by β2a subunits.

### Requirements for CaMKII-dependent Ca_V_1.3 complex assembly

To determine whether both Ca^2+^ and calmodulin are required for CaMKIIα-dependent assembly of multimeric Ca_V_1.3 complexes, we tested whether Ca^2+^ alone or calmodulin alone would support complex formation in HEK cell lysates using the co-immunoprecipitation assay.

Negligible levels of mCherry-Ca_V_1.3 associated with iHA-Ca_V_1.3 in the absence of CaMKIIα under any conditions. Moreover, in contrast to the effect of adding both Ca^2+^ and calmodulin in the presence of CaMKIIα, neither Ca^2+^ nor calmodulin alone enhanced the co-immunoprecipitation of mCherry-Ca_V_1.3 with iHA-Ca_V_1.3 (Figure 2C-D). A similar result was obtained in the presence of FLAG-β3 (Figure S4A-B). In addition, calmidazolium (50 μM), a widely used calmodulin antagonist, essentially abrogated the Ca^2+^/calmodulin-dependent association of mCherry-Ca_V_1.3 and CaMKIIα with iHA-Ca_V_1.3 (Figure S5A-B). Taken together, these data indicate that both Ca^2+^ and calmodulin are required for activated CaMKIIα-dependent assembly of multimeric Ca_V_1.3 complexes.

### CaMKII-dependent assembly of Ca_V_1.3/Ca_V_1.2 complexes

Since the CaMKII-binding RKR motif is conserved between the cytosolic N-terminal domains of Ca_V_1.2 and Ca_V_1.3 (Wang *et al*., 2017b). Therefore, we tested the hypothesis that CaMKII can assemble protein complexes containing both Ca_V_1.2 and Ca_V_1.3 by HA-immunoprecipitation from lysates of HEK293T cells co-expressing iHA-Ca_V_1.3, GFP-Ca_V_1.2, α2δ, and FLAG-β2a with or without wild type CaMKIIα under basal conditions (with EDTA) or following Ca^2+^/calmodulin addition (Figure 2E). Under basal conditions, minimal levels of GFP-Ca_V_1.2 were detected in HA immune complexes in with or without co-expressed CaMKIIα. However, Ca^2+^/calmodulin addition to lysates containing CaMKIIα significantly increased the levels of GFP-Ca_V_1.2, CaMKIIα and FLAG-β2a in the iHA-Ca_V_1.3 complex by about ∼5-fold, significantly more than a modest ∼1.5 fold increase detected in the absence of co-expressed CaMKIIα (Figure 2F). In a second series of experiments, we found that the CaMKIIα-V102E mutation, which disrupts CaMKIIα binding to the N-terminal domains of both Ca_V_1.3 and Ca_V_1.2 (Figure S6), failed to support Ca^2+^/calmodulin-dependent assembly of Ca_V_1.2/Ca_V_1.3 complexes (Figure 2G-H). Taken together, these data support the hypothesis that direct interactions of N-terminal domains of α1 subunits with activated catalytic domains in CaMKIIα holoenzymes can assemble complexes containing both Ca_V_1.2 and Ca_V_1.3 α1 subunits *in vitro*.

### CaMKII-dependent assembly of multi-Ca_V_1.3 complexes in the presence of Shank3

Shank3 facilitates the assembly of multimeric Ca_V_1.3 complexes under basal conditions, but not following the addition of Ca^2+^/calmodulin (Yang *et al*., 2023), yet it also binds activated CaMKII (Perfitt *et al*., 2020). Therefore, we investigated the impact of Shank3 on Ca^2+^/calmodulin- and CaMKII-dependent assembly of Ca_V_1.3 complexes. Lysates of HEK293T cells co-expressing iHA-Ca_V_1.3, mCherry-Ca_V_1.3, α2δ, and FLAG-β2a with CaMKIIα or GFP-Shank3, or both CaMKIIα and GFP-Shank3, were immunoprecipitated using HA antibodies under basal conditions (EDTA) or following Ca^2+^/calmodulin addition (Figure 3A). In the absence of CaMKIIα, Shank3 associated with iHA-Ca_V_1.3, enhancing the co-immunoprecipitation of mCherry-Ca_V_1.3 α1 subunits under basal (EDTA) conditions, but Ca^2+^/calmodulin addition disrupted these interactions, as seen previously (Yang *et al*., 2023). Conversely, in the absence of Shank3, CaMKIIα co-expression enhanced the association of mCherry-Ca_V_1.3 and FLAG-β2a with iHA-Ca_V_1.3 only after Ca^2+^/calmodulin addition, as seen in Figure 2A-D. However, in the presence of both CaMKIIα and GFP-Shank3, substantially more mCherry-Ca_V_1.3 co-immunoprecipitated with iHA-Ca_V_1.3 from basal (EDTA) cell lysates, even though there were no significant changes in the levels of co-precipitated iHA-Ca_V_1.3 or GFP-Shank3 (relative to lysates lacking CaMKIIα), and no significant co-precipitation of inactive CaMKIIα (Figure 3B). Addition of Ca^2+^/calmodulin to cell lysates containing CaMKIIα and GFP-Shank3 significantly increased the recruitment of CaMKIIα into the iHA-Ca_V_1.3 complex relative to the absence of GFP-Shank3, in parallel with an increased precipitation of iHA-Ca_V_1.3 itself, but there was no further increase in the levels of co-precipitated mCherry-Ca_V_1.3. Moreover, in the presence of activated CaMKIIα, GFP-Shank3 was retained in iHA-Ca_V_1.3 complexes following Ca^2+^/calmodulin addition. Notably, the ratio of mCherry-Ca_V_1.3 to iHA-Ca_V_1.3 was significantly higher under both EDTA and Ca^2+^/calmodulin conditions in the presence of both GFP-Shank3 and CaMKIIα than it was under basal conditions with GFP-Shank3 alone, or Ca^2+^/calmodulin conditions with CaMKIIα alone (Figure 3A-B). These findings indicate that, in the presence of both Shank3 and CaMKIIα, Shank3 is predominantly responsible for assembly of multi-Ca_V_1.3 complexes under basal conditions, but that activation of CaMKIIα following Ca^2+^/calmodulin addition results in retention of Shank3 in the complex, further enhancing formation of multimeric Ca_V_1.3 complexes.

**Figure 3.**
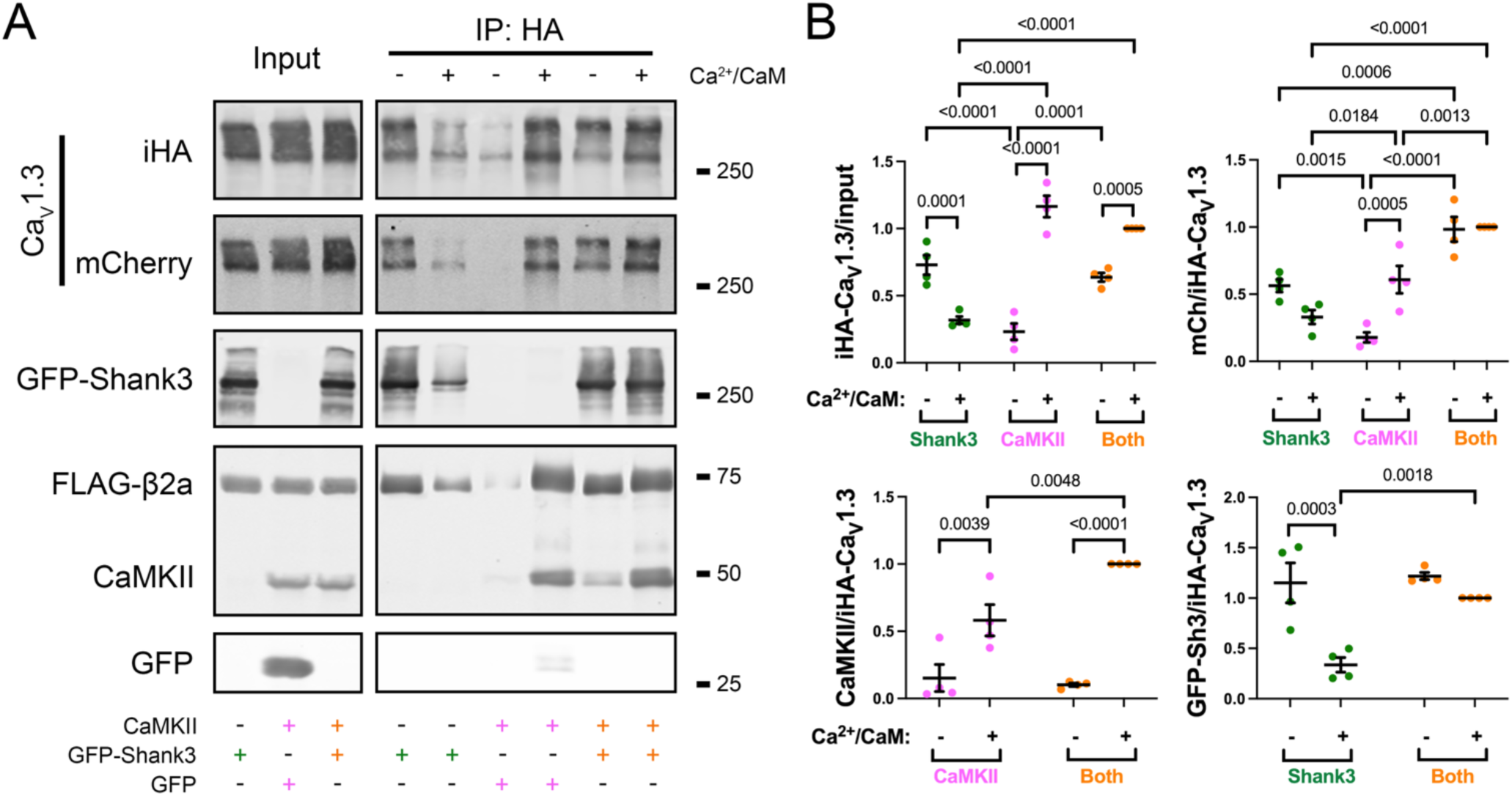
CaMKII-dependent assembly of Ca_V_1.3/β2a complexes prevents Ca^2+^/calmodulin-dependent dissociation of Shank3 when both proteins are present, enhancing complex formation. A) Representative immunoblots of the input and anti-HA IPs from soluble fractions of HEK293T cells co-expressing iHA- and mCherry-Ca_V_1.3, α2δ, and FLAG-β2a subunits, along with GFP-Shank3/pcDNA or GFP/CaMKII or GFP-Shank3/CaMKII. Half of each lysate was supplemented with, Ca^2+^/calmodulin prior to immunoprecipitation. Images of the immunoblots for each protein were from the same transfection replicate gel/blot and were collected using identical detection sensitivity. B) Quantification of iHA- and mCherry-Ca_V_1.3, CaMKII, and GFP-Shank3 signals in HA-immune complexes from four independent transfection replicates. Immunoblot signals were normalized as described in the legend to figure 2, relative to the “Ca^2+^/calmodulin + GFP-Shank3 + CaMKII” condition within each replicate. Statistical analyses: two-way ANOVA followed by Šídák’s post hoc tests (for comparisons without and with Ca^2+^/calmodulin condition) or by Tukey’s post hoc tests (for comparisons of iHA and mCh signals between different transfection conditions).

### CaMKII-dependent assembly of multi-LTCC complexes is disrupted by 1,6-hexanediol

To investigate the relationship between the *in vitro* assembly of multi-LTCC complexes by activated CaMKII and LTCC clustering in neurons, we compared the effects of 1,6 and 2,5-hexanediol on co-immunoprecipitation of multi-Ca_V_1.3 complexes in the presence of activated CaMKIIα. Lysates of HEK293T cells co-expressing iHA-Ca_V_1.3, mCherry-Ca_V_1.3, α2δ, FLAG-β2a, and CaMKIIα were supplemented with Ca²⁺/calmodulin (to activate CaMKIIα) alone, or in the additional presence of 1,6-hexanediol or 2,5-hexanediol (both at 2.5%). Notably, 1,6-hexanediol significantly decreased the co-immunoprecipitation of activated CaMKIIα with iHA-Ca_V_1.3 and reduced the association of mCherry-Ca_V_1.3 and FLAG-β2a with these complexes (Figure 4A-B). However, 2,5-hexanediol had no significant effect on the co-precipitation of these proteins.

**Figure 4.**
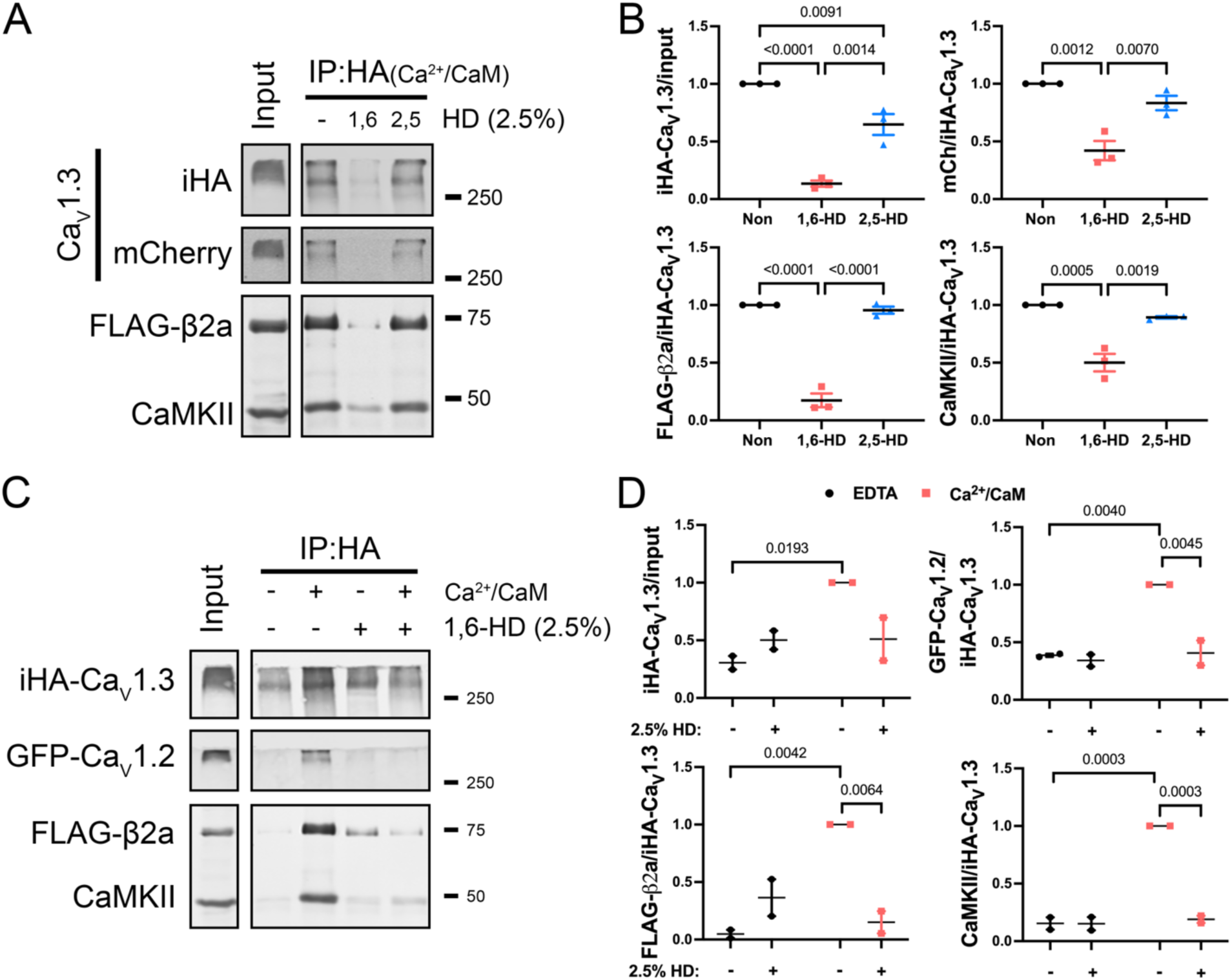
1,6-HD disrupts CaMKIIα- and Ca^2+^/calmodulin-dependent assembly of LTCC complexes in HEK293T cell lysates. A) Representative immunoblots in the input and anti-HA IPs from soluble fractions of HEK293T cells co-expressing iHA-Ca_V_1.3, mCherry-Ca_V_1.3, α2δ, and FLAG-β2a subunits, with CaMKII. Aliquots of the same lysates were supplemented with Ca^2+^ and calmodulin and the addition of vehicle (lysis buffer), 2.5% 1,6- or 2,5-hexanediol (HD). B) Quantification of iHA-Ca_V_1.3, mCherry-Ca_V_1.3, FLAG-β2a, and CaMKII signals in HA-immune complexes from three independent transfection replicates. Immunoblot signals were normalized as described in the legend to figure 2, relative to the Ca^2+^/calmodulin without HD condition within each replicate. Statistical analyses: one-way ANOVA with Tukey’s post hoc tests. C) Representative immunoblots of the input and anti-HA IPs from soluble fractions of HEK293T cells co-expressing iHA-Ca_V_1.3, GFP-Ca_V_1.2, α2δ, FLAG-β2a subunits, and CaMKIIα, without or with Ca^2+^/calmodulin addition in the absence or presence of 1,6-HD. D) Quantification of iHA-Ca_V_1.3, GFP-Ca_V_1.2, FLAG-β2a, and CaMKIIα signals in HA-immune complexes from two independent replicates. Immunoblot signals were normalized as described in the legend to figure 2, relative to the Ca^2+^/calmodulin condition without HD application within each replicate. Statistical analyses: two-way ANOVA with Šídák’s post hoc tests. Images of immunoblots for each protein in panels A and C were from the same transfection replicate gel/blot and were collected using identical detection sensitivity.

Similarly, we found that 1,6-hexanediol disrupted the activated CaMKIIα-dependent assembly of GFP-Ca_V_1.2/iHA-Ca_V_1.3 complexes, although there was no effect of 1,6-hexanediol under basal conditions (Figure 4C-D). In addition, we found that 1,6-hexanediol did not disrupt Shank3 dimerization, or the Shank3-dependent assembly of multi-Ca_V_1.3 complexes under basal conditions (Figure S7A-D). However, the association of activated CaMKIIα with Shank3 was significantly reduced by 1,6-hexanediol (Figure S7E-F). Taken together, these data support the hypothesis that the activated CaMKIIα-dependent assembly of multi-LTCC complexes *in vitro* involves some form of biomolecular condensate.

### CaMKIIα and Ca^2+^ influx enhances plasma membrane Ca_V_1.3 clustering in intact HEK293 cells

We next tested the hypothesis that activated CaMKII enhances the clustering of cell surface Ca_V_1.3 LTCCs in intact HEK293 cells. We co-expressed iHA-Ca_V_1.3, α2δ, and FLAG-β2a subunits with or without wild type CaMKIIα. Transfected cells were pre-incubated in a Ca^2+^-free HEPES buffer for 10 minutes and then in HEPES buffer with or without BayK (10 μM), an LTCC agonist, in the presence or absence of 2.5 mM Ca^2+^ for 5-10 minutes, prior to fixation/permeabilization. After immunolabeling for HA and CaMKIIα, we used total internal reflection fluorescence (TIRF) microscopy to detect immunofluorescent labels residing within ∼100 nm of the cover slip. As seen previously (Yang *et al*., 2023), iHA-Ca_V_1.3 was readily detected on, or near, the cell surface under all experimental conditions (Figure 5A), and we quantified the intensity and density of the puncta. In the absence of co-expressed CaMKIIα, the addition of Ca^2+^ and/or BayK had no significant effect on iHA-Ca_V_1.3 puncta intensity (Figure 5B-B’). Co-expression of CaMKIIα significantly, if modestly, increased puncta intensity in the absence of Ca^2+^ and BayK, and the addition of both Ca^2+^ and BayK significantly increased iHA-Ca_V_1.3 puncta intensity. Moreover, CaMKIIα co-expression modestly reduced iHA-Ca_V_1.3 cluster density independent of the cell treatment conditions (Figure 5C-C’). These data indicate that Ca^2+^ influx via Ca_V_1.3 LTCCs induces Ca_V_1.3 α1 subunit clustering in the plasma membrane of intact HEK293 cells via a CaMKII-dependent mechanism.

**Figure 5.**
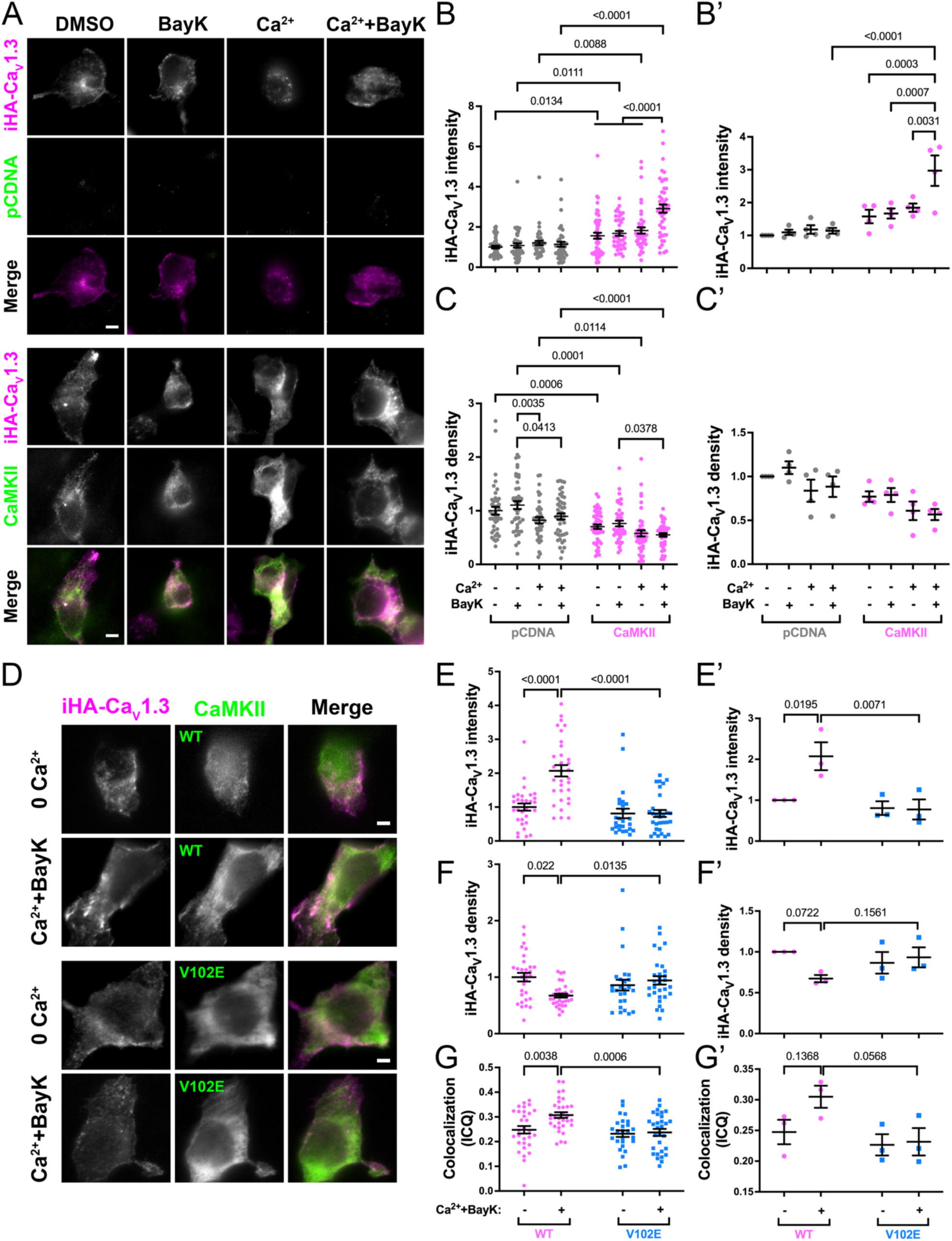
Activity- and CaMKIIα-dependent clustering of Ca_V_1.3/β2a LTCCs in HEK293 cells. A) Representative iHA-Ca_V_1.3, CaMKIIα and merged TIRF microscope images of fixed HEK293 cells co-expressing iHA-Ca_V_1.3, α2δ, and FLAG-β2a subunits, and either empty pCDNA vector or CaMKIIα. Cells were fixed after 5-10 minutes incubation with either no Ca^2+^ or Ca^2+^ buffer with vehicle DMSO or BayK 8644 (BayK, 10 μM), as indicated (scale bar, 5 μm). B-C) Quantification of iHA-Ca_V_1.3 puncta intensity (panel B) and density (panel C) from 8-13 cells per condition from 4 independent transfection (pCDNA: n = 43 for DMSO, 43 for BayK, 40 for Ca^2+^+DMSO, 45 for Ca^2+^+BayK; CaMKII: n = 50 for DMSO, 42 for BayK, 45 for Ca^2+^+DMSO, 47 for Ca^2+^+BayK). Data within each replicate were normalized to the mean of values in the absence of Ca^2+^ and BayK 8644. B’-C’) Re-analysis of data in panel B and C by the 4 replicates. Statistical analyses in B-C and B’-C’: two-way ANOVA followed by Šídák’s (for comparisons between pCDNA and CaMKII) or Tukey’s (for comparisons between different conditions) post hoc tests. D) Representative TIRF images of iHA-Ca_V_1.3 and CaMKII in fixed HEK293 cells co-expressing iHA-Ca_V_1.3, α2δ, and FLAG-β2a subunits, with wildtype CaMKII (WT) or CaMKII V102E mutant (V102E). Cells were incubated for approximately 5 min with zero Ca^2+^(0 Ca^2+^) or 2.5 mM Ca^2+^ and 10 μM BayK 8644 (Ca^2+^+BayK) prior to fixation. Scale bar, 5 μm. E-G) Quantification of iHA-Ca_V_1.3 puncta intensity (B) and density (C), as well as colocalization between iHA-Ca_V_1.3 and CaMKII (D), respectively from three independent transfections. Data derived from n = 30 (0 Ca^2+^) and 32 (Ca^2+^+BayK) cells expressing WT CaMKII, and n = 26 (0 Ca^2+^) and 30 (Ca^2+^+BayK) cells expressing V102E CaMKII. The iHA puncta staining intensity and density values were normalized to the mean WT value in the absence of Ca^2+^ and BayK 8644 within each replicate. E’-G’) Re-analysis of data in panel B-D by the 3 replicates. Statistical analyses in E-G and E’-G’: two-way ANOVA with Šídák’s post hoc tests.

To investigate the molecular basis of the CaMKII- and Ca^2+^/BayK-dependent increases in plasma membrane iHA-Ca_V_1.3 puncta intensity, we co-expressed iHA-Ca_V_1.3 LTCCs with either WT CaMKIIα (CaMKII-WT) or a V102E CaMKIIα mutant (CaMKII-V102E), which disrupts binding to the N-terminal domain of Ca_V_1.3/1.2 α1 subunits (Wang *et al*., 2017b). HEK293 cells were treated with either Ca^2+^-free HEPES buffer (0 Ca^2+^) or HEPES buffer supplemented with 2.5 mM Ca^2+^ and BayK (Figure 5D). In the presence of CaMKII-WT the intensity of surface-localized iHA-Ca_V_1.3 clusters was significantly increased following Ca^2+^/BayK addition, in parallel with a decrease in cluster density (Figures 5E-E’ and 5F-F’). However, Ca^2+^/BayK-induced changes in iHA-Ca_V_1.3 clustering were not observed in cells expressing the CaMKII-V102E mutant.

Concurrently, we assessed CaMKIIα colocalization with iHA-Ca_V_1.3 using the ICQ method (Figures 5G-G’). Under basal conditions, ICQ scores for cells co-expressing CaMKII-WT (∼0.25) and CaMKII-V102E (∼0.23) were similar. However, the Ca^2+^/BayK incubation significantly increased the ICQ score in cells co-expressing CaMKII-WT (∼0.31), but not in cells co-expressing CaMKII-V102E. Taken together, these findings indicate that binding of activated CaMKII to the Ca_V_1.3 N-terminal domain is required for Ca_V_1.3 clustering in the plasma membrane of HEK293 cells.

### Activity-dependent clustering of LTCCs in hippocampal neurons requires endogenous CaMKII

To test the role of CaMKII in neuronal LTCC clustering, we transfected hippocampal neurons at DIV14 to express Ca_V_1.3 or Ca_V_1.2 tagged with an extracellular HA epitope (sHA-Ca_V_1.3, sHA-Ca_V_1.2) and FLAG-β2a with either a nonsense shRNA (nssh) or specific shRNAs targeting CaMKIIα and CaMKIIβ (CaMKII-sh) (both with a soluble GFP marker). To establish the efficacy of the shRNA approach under these conditions, neurons were immunostained for CaMKIIα.

Neurons expressing control nssh contained somatic and dendritic pools of CaMKIIα, with no significant change in intensity compared to adjacent non-transfected neurons (Figure S8A-B). In contrast, the intensity of CaMKIIα staining in neurons expressing the CaMKII-sh was significantly reduced by about 77% relative to nearby non-transfected neurons (Figure S8B-B’). Thus, the shRNAs effectively suppressed CaMKII expression under the current experimental conditions.

To confirm the biological role of CaMKII when sHA-Ca_V_1.3 is over-expressed in neurons, we depolarized neurons using 40mM KCl for 90 seconds (as in Figure 1), fixed the neurons and then labeled surface-localized sHA-Ca_V_1.3. Neurons were then permeabilized and labeled for Ser133 phosphorylation of CREB (pCREB) (Figure S9A). Depolarization of neurons expressing sHA-Ca_V_1.3 with control nssh shRNA robustly increased CREB phosphorylation, and to a slightly, but significantly, greater extent compared to nearby non-transfected neurons (Figures S9B-B’).

However, depolarization of neurons expressing sHA-Ca_V_1.3 and the CaMKII-sh shRNA resulted in significantly ∼43% lower levels of nuclear CREB phosphorylation compared to nearby non-transfected neurons (Figures S9C-C’). Taken together, these data indicate that E-T coupling in DIV21 neurons overexpressing sHA-Ca_V_1.3 was partially CaMKII-dependent, as seen in DIV14 neurons expressing only endogenous LTCCs (Wheeler *et al*., 2008; Wang *et al*., 2017b).

Next, we collected super-resolution images of surface-expressed sHA-Ca_V_1.3 or sHA-Ca_V_1.2 LTCCs using the Airyscan mode of a Zeiss LSM880 microscope and quantified LTCC puncta intensity and density with or without neuronal depolarization (Wu and Hammer, 2021; Yang *et al*., 2023). In neurons expressing control nssh shRNA, depolarization significantly increased the intensity of both sHA-Ca_V_1.3 and sHA-Ca_V_1.2 puncta in the soma and dendrites compared to 5K treatment (data for dendritic and somatic analyses are shown in Figure 6 and Figure S10, respectively), consistent with changes observed in Figure 1. However, in neurons expressing CaMKII-sh shRNA, depolarization resulted in a significant decrease in sHA-Ca_V_1.3 puncta intensity in both the soma and dendrites (Figure 6A, 6C, S10A, and S10C), but no change in the intensity of dendritic sHA-Ca_V_1.2 puncta and a modest decrease in the intensity of somatic sHA-Ca_V_1.2 puncta (Figure 6D, 6F, S10D, and S10F). There was no significant effect of depolarization on the density of somatic or dendritic sHA-Ca_V_1.3 puncta with or without CaMKII knockdown (Figure 6B-C and S10B-C), but depolarization increased sHA-Ca_V_1.2 puncta density in the soma (Figure S10F), but not the dendrites (Figure 6F), of CaMKII-sh-expressing neurons. Figure S11 shows the same data quantified and compared by the number of independent replicates performed. Taken together, these data indicate that CaMKII is important for increased clustering of surface localized Ca_V_1.3 and Ca_V_1.2 LTCCs following a brief depolarization under conditions that initiate E-T coupling.

**Figure 6.**
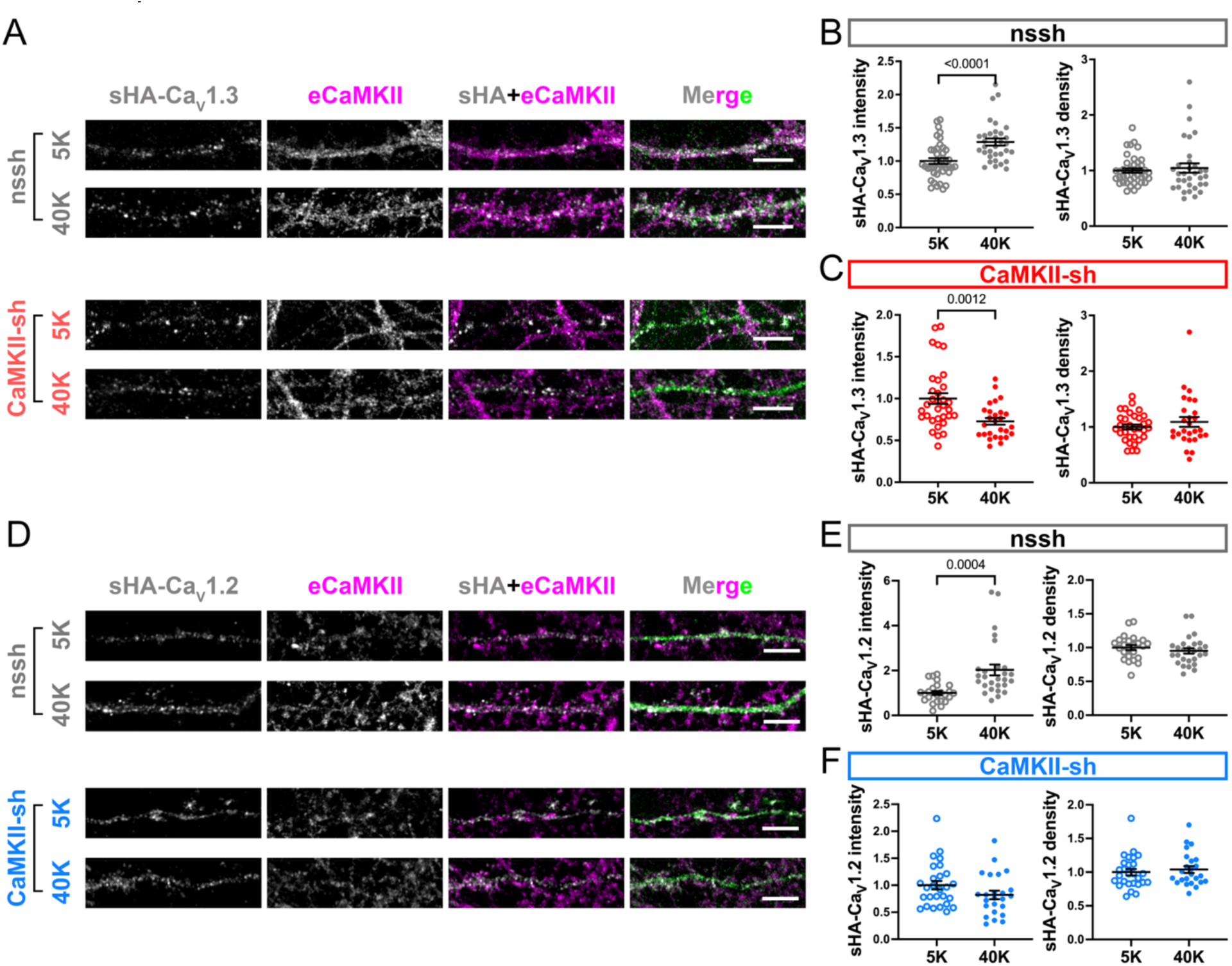
Depolarization-induced Ca_V_1.3 and Ca_V_1.2 clustering in hippocampal neurons is disrupted by CaMKII knockdown. Hippocampal neurons expressing sHA-Ca_V_1.3/sHA-Ca_V_1.2 and FLAG-β2a with either GFP-nssh or GFP-CaMKII-sh (DIV 21) were incubated for 90 s with 40K/5K and fixed (Figure 1). Neurons were immunostained for HA, permeabilized, and then endogenous CaMKII (eCaMKII) was immunostained. Images collected using Airyscan super-resolution confocal microscopy. A) Representative images of sHA-Ca_V_1.3 and CaMKII staining in dendrites of neurons expressing GFP-nssh. Scale bar, 5 μm. B-C) Quantification of sHA-Ca_V_1.3 cluster intensity and density in GFP-nssh (B) and GFP-CaMKII-sh (C) expressing neurons (n = 41 (5K), 33 (40K) neurons expressing nssh; n = 33 (5K) and 27 (40K) neurons expressing CaMKII-sh). Data were quantified from 5 independent cultures/transfections and normalized to the mean for transfected neurons following the 5K incubation within each replicate. Statistical analyses: unpaired t-tests. D) Representative images of sHA-Ca_V_1.2 and CaMKII staining in dendrites of neurons expressing GFP-CaMKII-sh. Scale bar, 5 μm. E-F) Quantification of sHA-Ca_V_1.2 cluster intensity and density in GFP-nssh (E) and GFP-CaMKII-sh (F) expressing neurons (n = 24 (5K), 27 (40K) neurons expressing nssh; n = 27 (5K) and 24 (40K) neurons expressing CaMKII-sh). Data were quantified from 3 independent cultures/transfections and normalized to the mean for transfected neurons following the 5K incubation within each replicate. Statistical analyses: unpaired t-tests. Supplementary figures 9 and 10 show quantitative analyses of images of the neuronal soma from these experiments.

We next examined co-clustering of Ca_V_1.3 and Ca_V_1.2 LTCCs in cultured hippocampal neurons. Neurons were co-transfected with sHA-Ca_V_1.3, FLAG-β2a, and either nssh or CaMKII-sh at DIV 14, and after 7 days (DIV21) they were depolarized with 40mM KCl for 90 seconds, fixed, and then immunostained for sHA-Ca_V_1.3 (without permeabilization) and endogenous Ca_V_1.2 (following permeabilization) (Figure 7A and S12A). Regions of interest (ROIs) were defined based on the sHA-Ca_V_1.3 signal and the amount of endogenous Ca_V_1.2 staining within each ROI was quantified separately in the dendrites (Figures 7B-B’) and soma (Figures S12B-B’). This approach allowed us to quantify Ca_V_1.2 signals derived predominantly from transfected neurons and not nearby non-transfected cells, and the resulting intensity ratio is proportional to the degree of co-localization between Ca_V_1.2 and sHA-Ca_V_1.3 on the cell surface. We found that the eCa_V_1.2/sHA-Ca_V_1.3 staining ratio was relatively low in both the soma and dendrites under basal (5K treatment) conditions (mean normalized to 1.0) and was unaffected by CaMKII knockdown. However, depolarization resulted in a significant increase in the eCa_V_1.2/sHA-Ca_V_1.3 staining ratio in the soma and dendrites of neurons expressing control shRNAs but not following CaMKII knockdown. Since depolarization increased the clustering of both Ca_V_1.3 and Ca_V_1.2 individually, we interpret the depolarization-induced increase of the eCa_V_1.2/sHA-Ca_V_1.3 staining ratio as representing increased co-clustering of Ca_V_1.2 with sHA-Ca_V_1.3, analogous to the co-immunoprecipitation of Ca_V_1.2 with Ca_V_1.3 in the presence of activated CaMKIIα (Figure 2E-H). Closer qualitative examination of images indicated that a subfraction of endogenous Ca_V_1.2 appeared to be localized to dendritic spines, whereas most of the sHA-Ca_V_1.3 staining, and the co-clustered channels appear to be localized to dendritic shafts (Figure 7A). Overall, these data demonstrate that neuronal depolarization induces the plasma membrane co-clustering of Ca_V_1.2 with Ca_V_1.3 via a CaMKII-dependent mechanism.

**Figure 7.**
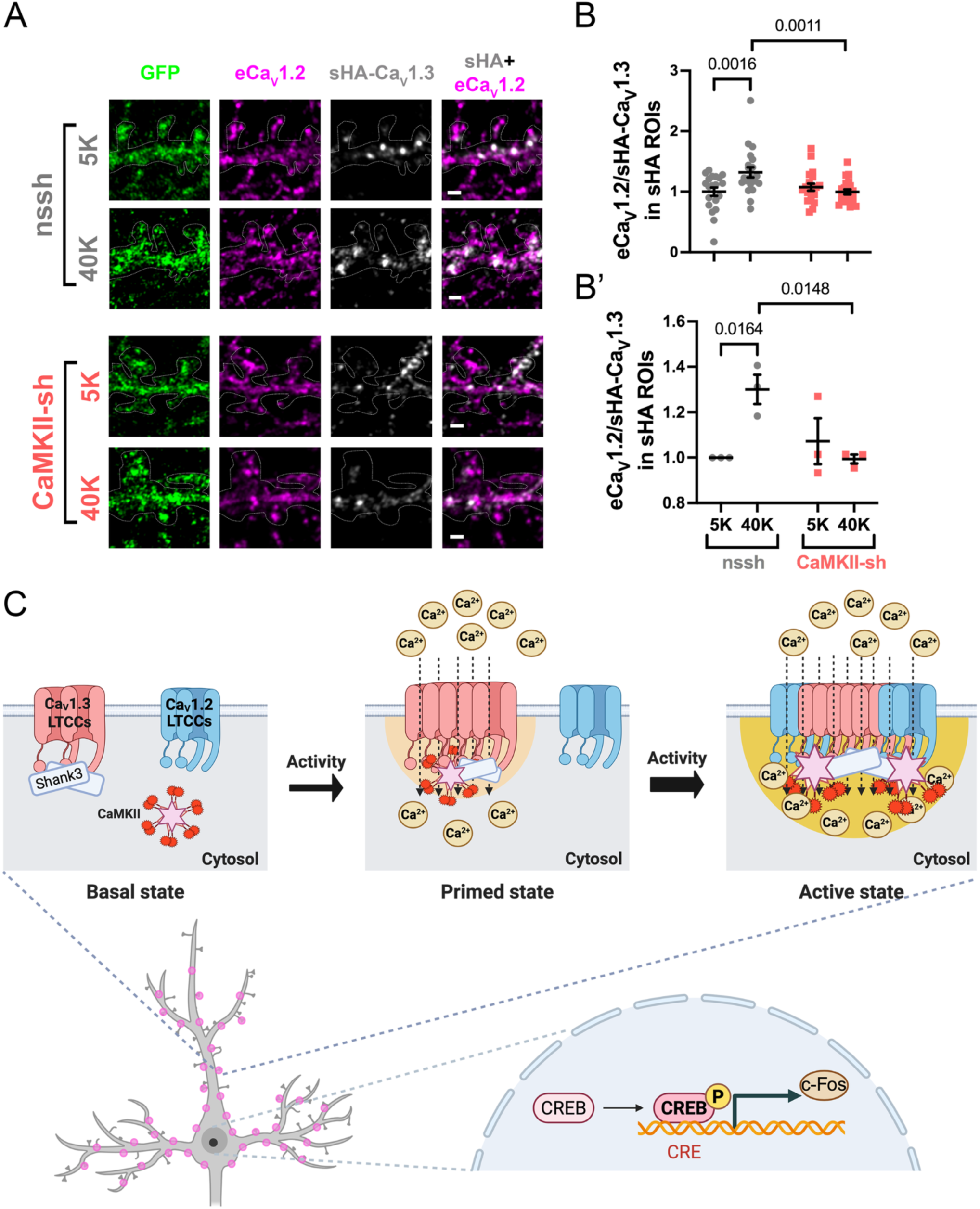
Depolarization-induced Ca_V_1.3-Ca_V_1.2 co-clustering is CaMKII-dependent in hippocampal neurons. Hippocampal neurons expressing sHA-Ca_V_1.3 and FLAG-β2a with either GFP-nssh or GFP-CaMKII-sh (DIV21) were incubated for 90 s with 40K/5K and fixed (see Figure 1). Neurons were immunostained for HA, permeabilized and then immunostained for endogenous Ca_V_1.2 (eCa_V_1.2). Neurons were imaged using an Airyscan super-resolution confocal microscopy. A) Representative images (2 per condition) of sHA staining, eCa_V_1.2 staining, and merged images in dendrites of neurons expressing GFP-nssh or CaMKII-sh. Scale bar, 5 μm. B) ROIs were defined based on sHA staining in order to measure the ratio of normalized eCa_V_1.2 intensity to normalized sHA intensity. Quantification of eCa_V_1.2 staining within sHA-Ca_V_1.3 ROIs (see methods) in neurons from three independent transfected cultures. n = 20 (5K) and 21 (40K) neurons expressing nssh; n = 20 (5K) and 22 (40K) neurons expressing CaMKII-sh. Data were normalized to the average eCa_V_1.2/sHA-Ca_V_1.3 ratio in nssh neurons with 5K treatment for each transfection. B’) Re-analysis of data in panel B by the 3 replicates. Statistical analyses: two-way ANOVA with Šídák’s post hoc tests. C) Proposed model for neuronal activity induced co-clustering of Ca_V_1.3 and Ca_V_1.2 LTCCs via CaMKII, forming Ca²⁺ nanodomains that initiate signaling to the nucleus. **Top**: Schematic of molecular events at the membrane. In the basal state, Ca_V_1.3 and Ca_V_1.2 exist in separate clusters, with Ca_V_1.3 clusters formed by Shank3 and CaMKII in the cytosol. Modest depolarization preferentially activates Ca_V_1.3 leading to Ca²⁺ influx (light yellow gradient) and CaMKII activation, triggering CaMKII recruitment, retention of Shank3 and enlargement of the Ca_V_1.3 clusters, via phase separation, forming a primed state. Sustained CaMKII activation and Ca_V_1.3 clustering leads to Ca_V_1.2 recruitment, forming larger Ca_V_1.2/Ca_V_1.3 co-clusters that amplify the Ca²⁺ nanodomain (dark yellow gradient). **Bottom**: Cartoon of an activated hippocampal neuron with CaMKII-mediated Ca_V_1.2/Ca_V_1.3 co-clusters and associated Ca^2+^ nanodomains distributed throughout the dendrites and soma (purple dots), which initiate signaling to the nucleus, represented by phosphorylation of CREB to drive transcription of immediate early genes such as *c-Fos*.

## Discussion

Our findings provide new insights into molecular mechanisms that control clustering of the neuronal LTCCs that are major initiators of E-T coupling, a key cellular process underlying synaptic plasticity, learning and memory (Ma *et al*., 2023). We initially found that the clustering of both Ca_V_1.2 and Ca_V_1.3 LTCCs in hippocampal neurons is increased following a brief depolarization that initiates E-T coupling. Notably, 1,6-hexanediol selectively disrupts LTCC clustering, as well as E-T coupling, compared to 2,5-hexanediol, indicating that both LTCC clustering and E-T coupling involve some form of biomolecular condensation. Using co-immunoprecipitation and fluorescence microscopy experiments, we found that activated CaMKII holoenzymes assemble homo-or hetero-meric Ca_V_1.3 or Ca_V_1.2 LTCC complexes *in vitro* and in the plasma membrane of HEK cells. Multi-LTCC complex assembly is facilitated by co-expression of LTCC β2a subunits (relative to the β3 subunits) and by Shank3, both of which also bind to CaMKII, and is disrupted by mutations in the CaMKII catalytic domain (V102E) or the N-terminal domain of Ca_V_1.3 α1 subunits (Δ69-93) that prevent their interaction. Significantly, the *in vitro* assembly of multi-LTCC complexes is also selectively disrupted by 1,6-hexanediol.

Moreover, clustering and co-clustering of neuronal Ca_V_1.2 and Ca_V_1.3 channels is impaired by shRNA-mediated knockdown of CaMKII expression. Taken together, these findings indicate that neuronal depolarization initiates E-T coupling by inducing the CaMKII-dependent clustering of LTCCs in a biomolecular condensate (Figure 7B).

### Clustering of L-type Ca^2+^ channels

LTCCs have a propensity to cluster in the plasma membrane, in some cases with other ion channels, modulating action potentials in neurons and neuroendocrine cells (Vivas *et al*., 2017; Cox, 2014; Marcantoni *et al*., 2010; Vandael *et al*., 2010; Plante, Whitt and Meredith, 2021). For example, plasma membrane clustering of Ca_V_1.2 with voltage-gated K_V_2.1 potassium channels in the neuronal soma and proximal dendrites facilitates additional interactions with ryanodine receptors (RyRs) in the endoplasmic reticulum membrane, facilitating Ca^2+^ release from endoplasmic reticulum (Vierra *et al*., 2019). Moreover, disruption of Ca_V_1.2-K_V_2.1 clustering interferes with E-T coupling (Vierra *et al*., 2021). Other studies indicate that LTCCs can form clusters containing an average of approximately eight α1 subunits in both adult ventricular myocytes (Ca_V_1.2) and cultured hippocampal neurons (Ca_V_1.3) (Dixon *et al*., 2015; Moreno *et al*., 2016). Notably, this clustering facilitates functional coupling of Ca_V_1.2 or Ca_V_1.3_S_ (isoform with a short C-terminal domain that cannot bind to Shank3), enhancing Ca^2+^ influx following depolarization due to interactions of the two lobes of Ca^2+^/calmodulin with α1 subunit C-terminal domains (Fallon *et al*., 2009; Dixon *et al*., 2015; Moreno *et al*., 2016). In contrast, the Ca_V_1.3_L_ isoform used in the current studies, which contains the full-length C-terminal domain including the Ca^2+^/calmodulin-binding site, but can bind to Shank3, forms similar sized clusters but without functional coupling. However, the molecular basis for physical clustering of LTCCs remains poorly understood.

Several proteins that interact with LTCC intracellular domains have been implicated in clustering, particularly if these proteins contain a PDZ domain and dimerize or multimerize (see Introduction). The C-terminal PDZ binding domain of Ca_V_1.2 LTCC appears to be important for surface localization in cultured hippocampal neurons (Weick *et al*., 2003), while the C-terminal PDZ-binding domain of Ca_V_1.3_L_ interacts with the PDZ domain of Shank3, a synaptic scaffolding protein, to facilitate surface expression (Zhang *et al*., 2005; Zhang *et al*., 2006). These effects on surface expression appear to be relatively insensitive to depolarization and independent of CaMKII expression on the short (90 s) time scales analyzed in the current studies because we detected no consistent changes of LTCC cluster density in the soma or dendrites when data were analyzed across independent biological replicates (Figures S1, S11). However, Shank3 homo-multimerizes via interactions between C-terminal SAM domains (Sheng and Kim, 2000), and we recently reported that Shank3 clusters multiple Ca_V_1.3 LTCCs under basal (low Ca^2+^) conditions *in vitro*, in HEK293 cells, and in cultured hippocampal neurons (Yang *et al*., 2023).

However, Shank3 knockdown does not affect basal clustering of neuronal Ca_V_1.2 LTCCs, which cannot bind to the Shank3 PDZ domain. Notably, Ca^2+^ disrupts Shank3 dependent Ca_V_1.3 clustering *in vitro* and in HEK293 cells in parallel with the dissociation of Shank3 from the complex (Yang *et al*., 2023). However, our current findings show that neuronal depolarization enhances LTCC clustering. We will further discuss this discrepancy below.

### Role for biomolecular condensates in neuronal LTCC clustering following depolarization

One remarkable feature of LTCC-dependent E-T coupling is that it can be blocked by loading neurons with BAPTA to chelate intracellular Ca^2+^, but is preserved in neurons loaded EGTA (Deisseroth, Bito and Tsien, 1996; Ma *et al*., 2012). This observation indicates that E-T coupling can be initiated by Ca^2+^ signaling events in a nanodomain in the vicinity of LTCCs that can be blocked only by the fast Ca^2+^ chelator, BAPTA, without requiring global Ca^2+^ increases, that are blocked by both BAPTA and the slow Ca^2+^ chelator, EGTA. Subsequent studies showed that CaMKII recruitment to the LTCC Ca^2+^ signaling nanodomain is required to initiate E-T coupling (Wheeler *et al*., 2008; Ma *et al*., 2014; Wang *et al*., 2017b). It seems intuitive that the efficiency of E-T coupling will depend on the dynamic properties of these Ca^2+^ nanodomains (e.g., amplitude, duration and size), which may in turn be modulated by LTCC clustering. We found that a neuronal depolarization paradigm that initiates E-T coupling (i.e., phosphorylation of CREB) also enhances the clustering of both Ca_V_1.2 and Ca_V_1.3 LTCCs.

Biomolecular condensates have recently emerged as a mechanism for formation of membrane-less subcellular compartments in cells (Gao *et al*., 2022; McSwiggen *et al*., 2019; Musacchio, 2022). Extensive *in vitro* studies have shown that mixtures of purified domains from multiple postsynaptic scaffolding proteins can assemble in liquid-liquid phase separated (LLPS) droplets, particularly at high concentrations (Zeng *et al*., 2018). Furthermore, inactive CaMKII can form LLPS droplets with fragments of Shank3 (Cai *et al*., 2021), and it has been shown that activated CaMKII can partition into LLPS droplets preformed by mixing fragments of the GluN2B C-terminal domain, Shank3 and other postsynaptic proteins (Cai *et al*., 2023; Hosokawa *et al*., 2021). However, conclusive evidence supporting a physiological role for biomolecular condensates containing full-length postsynaptic proteins is largely lacking.

A recent paper showed that presynaptic voltage-gated Ca^2+^ channels participate in biomolecular condensates with other components of the synaptic vesicle release machinery (Jin *et al*., 2025). Moreover, hexanediols were recently used to implicate biomolecular condensation in the presynaptic control of evoked synaptic transmission, but not spontaneous transmission (Guzikowski and Kavalali, 2024), illustrating how the differential sensitivity to hexanediols can provide insights into physiological roles of biomolecular condensates.

Therefore, we explored a potential role for biomolecular condensates in postsynaptic LTCC clustering by comparing the effects of 1,6-and 2,5-hexanediols; the more widely spaced hydroxyl groups in 1,6-hexanediol favor selective disruption of some biomolecular condensates in comparison to 2,5-hexanediol (Kato, Zhou and McKnight, 2022). We found that 1,6-hexanediol, but not 2,5-hexanediol, present only during the 90 s depolarization, disrupted depolarization-induced increases in LTCC clustering in the neuronal plasma membrane, but had no effect on basal LTCC clustering (Figure 1D-F). Notably, since we directly depolarized the neurons using KCl in the presence of a sodium channel blocker and glutamate receptor antagonist, our findings are not confounded by diol effects on evoked synaptic transmission (Guzikowski and Kavalali, 2024). A caveat to these studies is that hexanediols can have other effects on cells that may be unrelated to disruption of biomolecular condensates, particularly at 5X higher concentrations (≥5% v/v) and for longer times (Düster *et al*., 2021; Meduri *et al*., 2022). However, a recent paper reported that phase-separated condensates of postsynaptic scaffolding proteins are insensitive to 10% 1,6-hexanediol (Jia *et al*., 2025). Thus, taken together these data suggest that LTCC clustering requires the formation of a 1,6-hexanediol-sensitive biomolecular condensate in response to neuronal depolarization.

### Activity-dependent formation of multi-LTCC complexes by CaMKII

CaMKII has been reported to interact with N- and C-terminal domains of the LTCC Ca_V_1.2 α1 subunit, modulating their biophysical properties or intracellular trafficking, respectively (Hudmon *et al*., 2005; Simms *et al*., 2015). We recently identified an additional CaMKII-binding “RKR” motif in the N-terminus of the Ca_V_1.3 α1 subunits that selectively binds activated CaMKII and showed that this interaction is required for neuronal LTCC-dependent E-T coupling (Wang *et al*., 2017b). Notably, the RKR motif is conserved in the N-terminal domain of Ca_V_1.2, but is not contained within the N-terminal CaMKII-binding domain identified by Simms et al (2015).

The current data provide new insights into the role of activated CaMKII binding to N-terminal domains of both Ca_V_1.2 and Ca_V_1.3, indicating that the interaction of multiple LTCCs with catalytic domains within a CaMKII holoenzyme mediates the formation of heteromultimeric Ca_V_1.2/Ca_V_1.3 complexes *in vitro*, LTCC clustering in the plasma membrane of intact transfected HEK293 cells, and enhanced LTCC clustering in response to neuronal depolarization.

### β2a subunit facilitate formation of CaMKII mediated multi-LTCC complexes

Four isoforms of the voltage-gated calcium channel β auxiliary subunits play unique roles in modulating the biophysical properties of the pore-forming α1 subunits as well as their trafficking to the plasma membrane (Buraei and Yang, 2010; Dolphin, 2003; Jarvis and Zamponi, 2007). We previously reported that activated CaMKII selectively binds to the β1 and β2a subunits, but not to β3 or β4 subunits (Grueter *et al*., 2008), enhancing CaMKII association with Ca_V_1.2 LTCCs (Abiria and Colbran, 2010). Furthermore, mutation of β2a to prevent CaMKII binding disrupts CaMKII-dependent facilitation of Ca_V_1.2 LTCCs in cardiomyocytes (Koval *et al*., 2010). The present biochemical studies demonstrated the Ca^2+^/calmodulin- and CaMKII-dependent assembly of multi-LTCC complexes is significantly enhanced by β2a relative to β3 (Figure 2). It will be interesting to investigate the potential roles of direct CaMKII-binding to β2a as well as of β2a palmitoylation (which enhances LTCC plasma membrane targeting (Gao, Chien and Hosey, 1999)) in LTCC (co-)clustering.

### Modulation of multi-LTCC complex formation by Shank3

Shank3 also directly interacts with both Ca_V_1.3 and activated CaMKII; disruption of either the Shank3-Ca_V_1.3 or the Shank3-CaMKII interaction interferes with LTCC-mediated E-T coupling in cultured hippocampal neurons (Perfitt *et al*., 2020). Given the role of Shank3 in basal Ca_V_1.3 clustering (see above), we were initially surprised that depolarization enhanced neuronal Ca_V_1.3 clustering leading us to uncover critical roles for CaMKII in depolarization-induced clustering of both Ca_V_1.3 and Ca_V_1.2 reported here. However, our *in vitro* experiments revealed that the formation of multimeric Ca_V_1.3 complexes under basal conditions is surprisingly enhanced by co-expression of CaMKIIα with Shank3 compared to the expression of either CaMKIIα or Shank3 alone (Figure 3), even though inactive CaMKIIα is not associated with Ca_V_1.3 complexes under basal conditions, in the presence or absence of Shank3. Further studies will be needed to determine how inactive CaMKII can enhance Shank3-dependent clustering under basal conditions; this may involve a reported interaction of inactive CaMKII with an N-terminal domain in Shank3 (Cai *et al*., 2021).

Ca^2+^/calmodulin addition to HEK293 cell lysates containing both CaMKIIα and Shank3 enhanced CaMKIIα association with iHA-Ca_V_1.3 complexes to a greater extent than observed when only CaMKIIα is present. Moreover, Shank3 remained associated with Ca_V_1.3 complexes following the addition of Ca^2+^/calmodulin in the presence of CaMKIIα, in contrast to the dissociation of Shank3 that was observed in the absence of CaMKIIα. We found that shRNA knockdown of neuronal CaMKII expression uncovered a significant depolarization-induced decrease in Ca_V_1.3 clustering (Figures 6A, 6C, S9A, and S9C), as might be predicted based on our prior *in vitro* and HEK293 studies of the Ca_V_1.3-Shank3 interaction. These observations raise further questions about the role of Shank3 in LTCC clustering following depolarization, especially in CaMKII-expressing neurons, because Shank3 cannot interact with the Ca_V_1.2 C-terminal domain. Taken together, these findings indicate that further studies will be required to more fully understand the complex interacting roles of Shank3 and CaMKII in modulating the formation of multimeric LTCC complexes following CaMKII activation.

### Role for biomolecular condensates in CaMKII-dependent formation of multi-LTCC complexes

As noted above, our initial data implicated biomolecular condensates in the depolarization-induced increase in LTCC clustering (see above). To test for a potential parallel role for biomolecular condensates in the *in vitro* formation of multi-LTCC complexes by activated CaMKII holoenzymes, we examined the sensitivity of these complexes to hexanediols. Our data indicate that 2.5% 1,6-hexanediol selectively disrupts the *in vitro* assembly of multi-Ca_V_1.2/1.3 complexes by activated CaMKII, whereas 2.5% 2,5-hexanediol had little effect. Significantly, the effects of 1,6-hexanediol appear to be specific for complexes involving activated CaMKII, since Shank3 dimerization and basal Shank3-dependent Ca_V_1.3 complex formation were insensitive to 1,6-hexanediols at these concentrations, whereas the co-immunoprecipitation of activated CaMKII with Shank3 was reduced by ∼75% by 1,6-hexanediol. These data indicate that activated CaMKII holoenzymes play a key role in the formation of molecular condensates involving these full-length proteins. This contrasts with prior studies showing that activated CaMKII partitions into LLPS droplets *in vitro* pre-formed from high concentrations of purified fragments of other postsynaptic proteins, in some cases re-organizing the protein fragments within the droplets (Hosokawa *et al*., 2021).

### LTCC-dependent E-T coupling is disrupted by 1,6-hexanediol

Commensurate with the disruption of neuronal LTCC clustering and the disruption of the formation of multi-LTCC complexes *in vitro*, we also found that 1,6-hexanediol selectively interfered with E-T coupling, measured by phosphorylation of nuclear CREB, even when present only during the 90 s depolarization. Notably, our use of CREB phosphorylation as a read out for E-T coupling mitigates concerns about potential direct effects of hexanediols on the actual transcriptional response (Düster *et al*., 2021; Meduri *et al*., 2022). Moreover, we found that cyclic AMP-induced CREB phosphorylation was insensitive to both of the hexanediols tested, suggesting that hexanediol disrupts upstream signaling pathway that are activated by neuronal depolarization. The fact that 1,6-hexanediol only partially blocked the increase in CREB phosphorylation is consistent with our studies showing that knockdown of CaMKII or Shank 3 expression only partially blocks CREB phosphorylation under these conditions (Figure S9), perhaps due the involvement of a non-CaMKII/Shank3 dependent pathway that is also independent of biomolecular condensation. Collectively, these data are consistent with the hypothesis that LTCC clustering in biomolecular condensates, driven by CaMKII activation, is an essential step for the efficient initiation of long-range E-T coupling mechanisms that signal to the nucleus.

### Role of activity-dependent Ca_V_1.2/1.3 co-clustering in E-T coupling

The highly homologous Ca_V_1.2 and Ca_V_1.3 LTCC α1 subunits are co-expressed in neurons at varying ratios across many brain regions where they appear to be preferentially localized to the soma, dendritic shafts, and dendritic spines (Striessnig *et al*., 2014; Folci *et al*., 2018; Stanika *et al*., 2016). This homology, and a lack of specific pharmacological reagents that can reliably modulate their activities, has made it challenging to identify unique physiological roles for Ca_V_1.2 and Ca_V_1.3, even though they have somewhat distinct biophysical properties. Perhaps most significantly, Ca_V_1.3 is more sensitive to membrane depolarization than Ca_V_1.2, presumably underlying previous observations that Ca_V_1.3 channels play a more dominant role in initiating neuronal E-T coupling in response to modest membrane depolarization (20-45 mM KCl), whereas the more abundant Ca_V_1.2 channels are dominant only with more robust depolarization (90 mM KCl) (Zhang *et al*., 2006). However, the combined use of transgenic mouse manipulations with LTCC agonists or blockers *in vivo* revealed important roles for both Ca_V_1.3 and Ca_V_1.2 in neuronal E-T coupling, with the relative contributions of the two channels varying between brain regions (Hetzenauer *et al*., 2006). The current findings suggest that CaMKII recruitment facilitates the co-clustering of Ca_V_1.3 and Ca_V_1.2 α1 subunits in the dendritic shaft and soma. These data indicate that Ca_V_1.3 and Ca_V_1.2 channels may “collaborate” to regulate downstream signaling under some circumstances. Since Ca_V_1.3 is preferentially activated by more modest membrane depolarization, we propose that the initial activation of Ca_V_1.3 and Ca^2+^ influx activates CaMKII, recruiting it to the channel to drive initial clustering of Ca_V_1.3. This might serve as a “priming” step to facilitate further recruitment of the more abundant (in hippocampal neurons) Ca_V_1.2 channels into a larger cluster (see Figure 8C) that may be required to create Ca^2+^ nanodomains with appropriate properties (e.g., size, amplitude, duration) for the initiation of E-T coupling. Furthermore, E-T coupling can also involve a “wave” of dendritic LTCC activation that is initiated in dendritic spines and propagates to the soma, apparently independent of sodium channel activity (Wild *et al*., 2019). We speculate that Ca_V_1.2/1.3 co-clusters, which appear to be predominantly present in dendritic shafts, may also be involved in this wave of dendritic LTCC activation. Further studies are required to test these ideas.

Taken together, our findings highlight a new layer of complexity in LTCC function—where dynamic, condensate-like channel assemblies enable synergistic interactions between channel subtypes, scaffold proteins, and signaling molecules. These assemblies may serve not only to organize channels spatially but also to modulate the precise characteristics of the resulting calcium signals and the amplitude and specificity of downstream signaling, such as CREB-mediated transcription. Further studies of the molecular determinants that regulate the formation, composition, and resolution of these CaMKII/LTCC condensates will be critical for elucidating their potential roles in both physiology and disease.

## Supporting information

PDF containing 12 supplementary figures

## Acknowledgements

This work was supported by Vanderbilt University and an endowed Louise B. McGavock Chair to R.J.C., and by an AHA fellowship award to X.W. (14PRE18420020). Imaging experiments and data analysis were performed in part through use of the Vanderbilt Cell Imaging Shared Resource (supported by National Institutes of Health Grants CA68485, DK20593, DK58404, DK59637, and EY08126). The content is solely the responsibility of the authors and does not necessarily represent the official views of the National Institutes of Health.

## Author contributions

Q.Y., X.W, and R.J.C designed research; Q.Y., X.W, and D.L. performed biochemistry experiments; Q.Y. performed imaging experiments; L.H. prepared rat hippocampal neuronal cultures; Q.Y. and L.H. performed neuron depolarization experiments; Q.Y. and D.L. analyzed data; drafts of the manuscript were originally written by Q.Y. and then edited by R.J.C. All authors participated in the discussion, revision, and approval of the final manuscript.

## Abbreviations

APV: D-2-amino-5-phosphonovalerate
BayK: BayK-8644
CaMKII: Calcium/calmodulin-dependent protein kinase II
CaMKII-sh: shRNAs targeting CaMKIIα and CaMKIIβ
CNQX: 6-Cyano-7-nitroquinoxaline-2,3-dione disodium
E-T: coupling excitation-transcription coupling
GFP: enhanced green fluorescent protein
GST: Glutathione-S-transferase
HA: Hemagglutinin
HD: hexanediol
ICQ: intensity correlation quotient
iHA: intracellular HA-tag
LLPS: liquid-liquid phase separation
LTCC: L-type calcium channel
Nssh: nonsense-shRNA
pCREB: Ser133-phosphorylated CREB
PDZ: domain PSD95/DlgA/Zo-1 domain
PSD: excitatory postsynaptic density
ROI: region of interest
SAM: Sterile alpha motif
sHA: extracellular HA tag
TIRF: Total internal reflection fluorescence
TTX: tetrodotoxin
WT: wild type

