## Supplementary material for "CaMKII clusters neuronal LTCCs in biomolecular condensates to gate excitation-transcription coupling": PDF containing 12 supplementary figures

**Supplemental Figures and legends**

**Supplemental Figure 1 (related to Figure 1). Depolarization-induced CaV1.2 clustering in hippocampal neurons is selectively disrupted by 1,6-hexanediol (HD).**
Re-analysis of data from Figure 1 (panels E, F, G) by averaging sHA-CaV1.2 cluster intensities and densities in soma (A) and in dendrites (B) within each independent replicate (n=3 cultures/transfections for neurons without HD or with 1% 1,6-HD, and 2 cultures/transfections for neurons with 1% 2,5-HD; 7-12 cells per transfection). Cluster densities and intensities within each replicate were normalized to the mean values in neurons following the 5K incubation without HD application. Statistical analyses: two-way ANOVA with Šídák's post hoc tests.

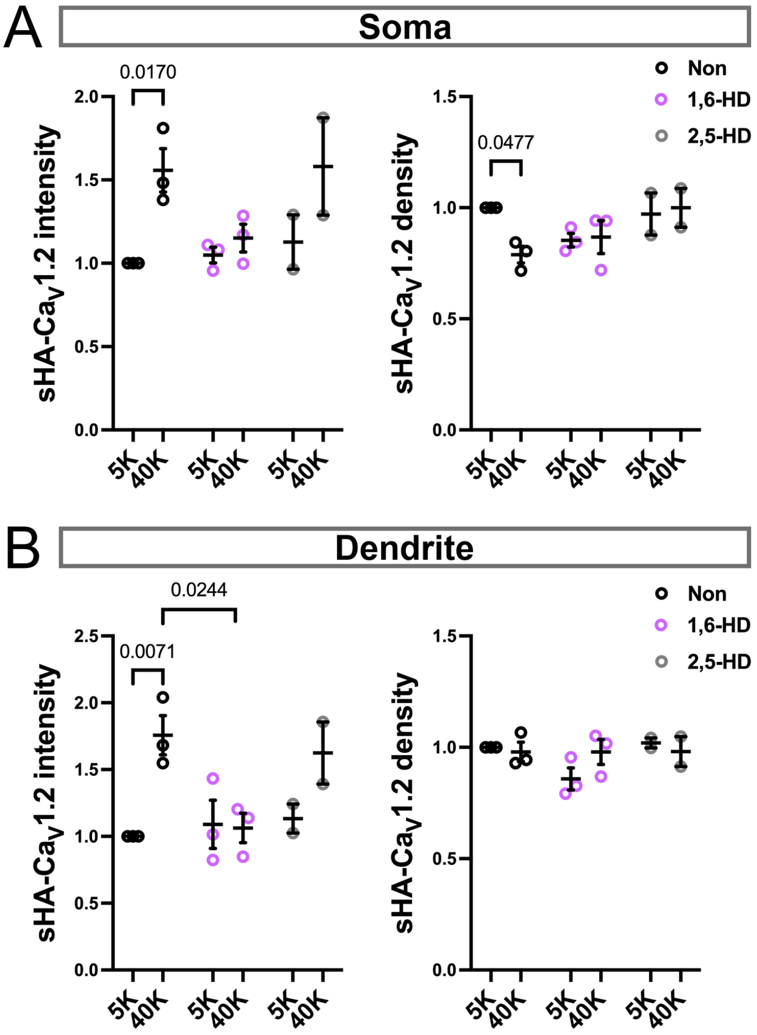

**Supplemental Figure 2 (related to Figure 1). Hexanediols do not affect forskolin stimulation of CREB phosphorylation at Serine 133.**
A) Representative images of pCREB, DAPI, and CaMKII immunostaining in hippocampal neurons (DIV 21) incubated for 5 min with or without 75 µM Forskolin and 1% 1,6- or 2,5-hexanediol (HD), as indicated. After fixation and staining, images were collected using confocal microscopy. Scale bar, 5 µm.
B) Quantification of nuclear pCREB signals in CaMKIIα-positive neurons (Non: n = 196; Forskolin: n =196; 1,6-HD: n = 271; 2,5-HD: n = 245).
B') Re-analysis of data in panel B by the 3 independent replicates.
Data were normalized to the mean for neurons following the Forskolin incubation without HD within each replicate. Statistical analyses: one-way ANOVA with Tukey's post hoc tests.

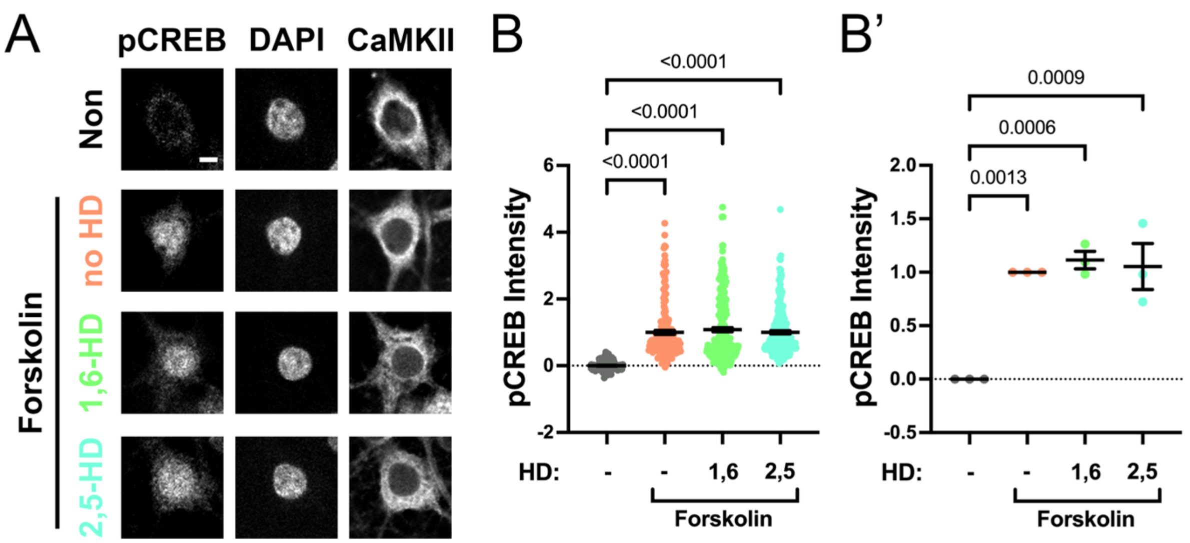

**Supplemental Figure 3 (related to Figure 2). Activated CaMKII holoenzymes assemble complexes containing multiple Cav1.3 α1 subunits.**

A. Immunoblots of the input and anti-HA IPs from soluble fractions of HEK293T cells co-expressing Cav1.3 (wild-type (WT) or NTD RKR motif deleted (Δ69-93)) with an N-terminal intracellular HA tag (iHA-Cav1.3), α2δ, and β3 subunits, with or without CaMKIIα (WT or association domain truncated (ΔAD)). Prior to immunoprecipitation, aliquots of lysates (containing EDTA) were supplemented with excess $Ca^{2+}$/calmodulin/$Mg^{2+}$-ATP.

B. Immunoblots of the input and anti-HA IPs from soluble fractions of HEK293T cells co-expressing iHA- (WT, Δ69-93 deleted or RKR mutated (AAA)) and FLAG-tagged Cav1.3, α2δ, and β3 subunits, with or without CaMKIIα (WT or V102E mutated). Prior to immunoprecipitation, aliquots of lysates (containing EDTA) were supplemented with excess $Ca^{2+}$/calmodulin/$Mg^{2+}$-ATP.

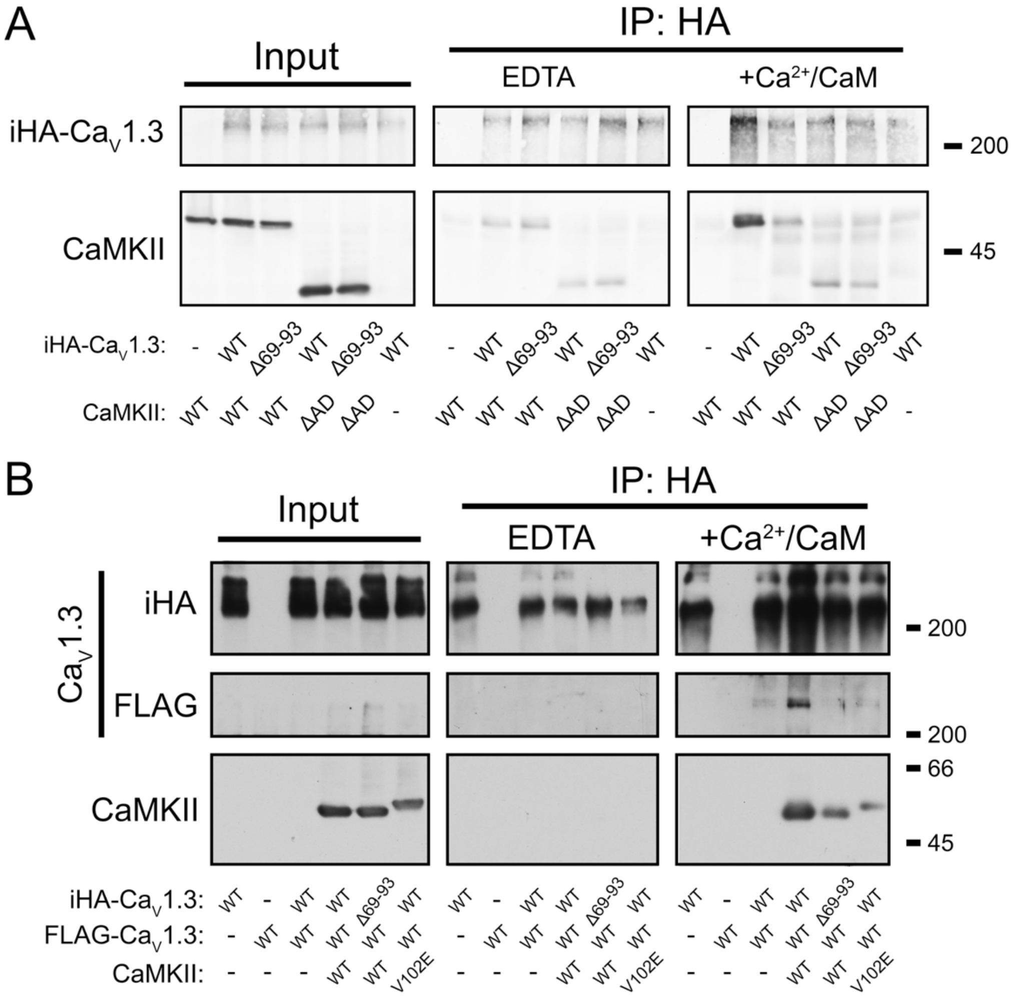

**Supplemental Figure 4 (related to Figure 2). CaMKII-dependent assembly of $Ca_V1.3$/β3 complexes is induced by $Ca^{2+}$/CaM.**

A) Representative immunoblots of the input and anti-HA IPs from soluble fractions of HEK293T cells co-expressing iHA-$Ca_V1.3$, mCherry-$Ca_V1.3$, α2δ, and FLAG-β3 subunits, with or without CaMKIIα. Prior to immunoprecipitation, aliquots of lysates were supplemented with $Ca^{2+}$ alone, calmodulin (CaM) alone, both $Ca^{2+}$ and calmodulin or an equal volume of $H_2O$. Images of the immunoblots for each protein were from the same transfection replicate gel/blot and were collected using identical detection sensitivity.

B) Quantification of iHA-$Ca_V1.3$ and mCherry-$Ca_V1.3$ signals in HA-immune complexes from three independent transfection replicates. iHA-$Ca_V1.3$ signals were normalized to the input signal, while mCherry-$Ca_V1.3$ signals were normalized to iHA-$Ca_V1.3$ in the corresponding IP. Protein signals within each replicate were then normalized to the $Ca^{2+}$/calmodulin/CaMKIIα condition to allow the replicate data to be pooled. Statistical analyses: two-way ANOVA with Šídák's (for comparisons without (black) and with (red) CaMKII co-expression) or Tukey's (for comparisons between four conditions) post hoc tests.

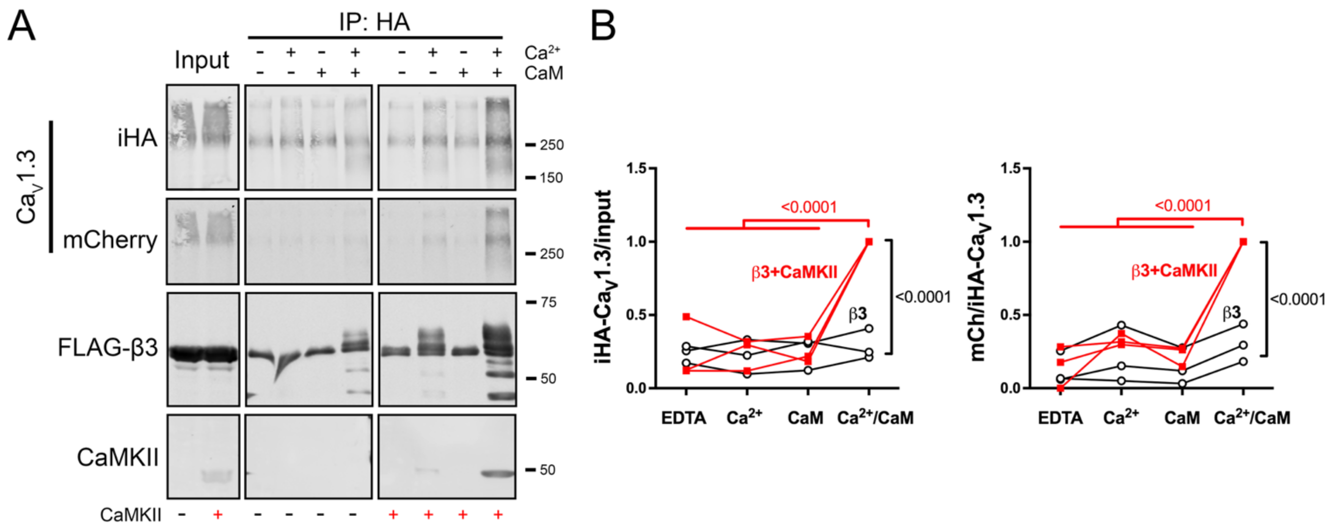

**Supplemental Figure 5 (related to Figure 2). Calmidazolium blocks CaMKII- and $Ca^{2+}$/calmodulin-dependent assembly of $Ca_V1.3$/β2a complexes in HEK 293T cell lysate.**
A) Representative immunoblots of the input and anti-HA IPs from soluble fractions of HEK293T cells co-expressing iHA-$Ca_V1.3$, mCherry-$Ca_V1.3$, α2δ, FLAG-β2a subunits and CaMKIIα, supplemented with or without added $Ca^{2+}$/calmodulin and with or without calmidazolium (50 μM), a calmodulin antagonist. Images of the immunoblots for each protein were from the same transfection replicate gel/blot and were collected using identical detection sensitivity.
B) Quantification of iHA-$Ca_V1.3$, mCherry-$Ca_V1.3$, and CaMKIIα signals in HA-immune complexes from three independent transfection replicates. iHA-$Ca_V1.3$ signals were normalized to the input signal; mCherry-$Ca_V1.3$ and CaMKIIα signals were normalized to iHA-$Ca_V1.3$ in the corresponding IP. Signals for each protein within each replicate were then normalized to the $Ca^{2+}$/calmodulin without calmidazolium condition to allow data from the replicates to be pooled. Statistical analyses: two-way ANOVA with Šídák's post hoc tests.

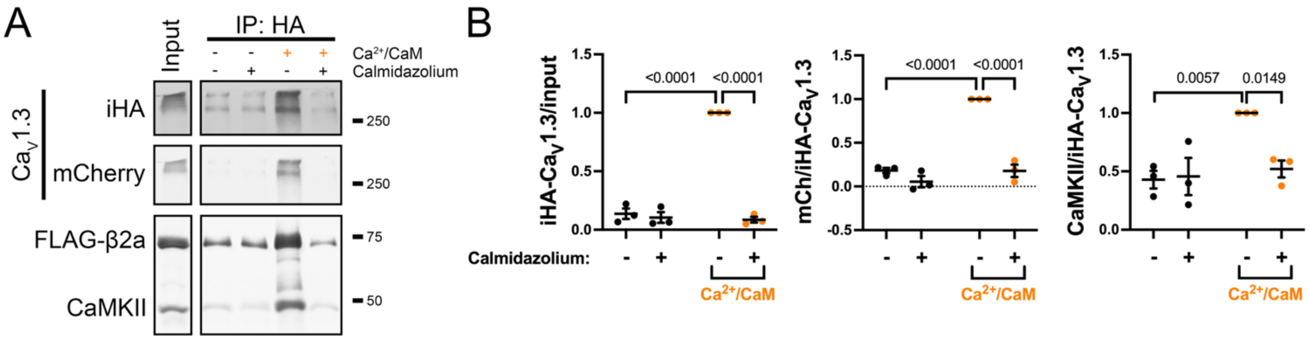

**Supplemental Figure 6 (related to Figure 2). CaMKIIα-V102E mutation disrupts association with both $Ca_V1.3$/β2a and $Ca_V1.2$/β2a.**

A) Representative immunoblots of the input and anti-HA IPs from soluble fractions from HEK293T cells co-expressing HA vector (-) or iHA-$Ca_V1.3$ (+), α2δ, FLAG-β2a subunits and CaMKIIα (WT or V102E). Equal aliquots of the lysates were supplemented with or without $Ca^{2+}$/calmodulin prior to immunoprecipitation.

B) Quantification of CaMKIIα and iHA-$Ca_V1.3$ signals in HA-immune complexes from three independent transfection replicates. iHA-$Ca_V1.3$ signals were normalized to the input signal, while CaMKII signals were normalized to iHA-$Ca_V1.3$ in the corresponding IP. Ratios within each replicate were normalized to the $Ca^{2+}$/calmodulin/CaMKIIα-WT condition to allow for pooling of the data. Statistical analysis: two-way ANOVA with Šídák's post hoc test.

C) Representative immunoblots of the input and anti-GFP IPs from soluble fractions of HEK293T cells co-expressing GFP vector (-) or GFP-$Ca_V1.2$ (+), α2δ, FLAG-β2a subunits, and CaMKIIα (WT or V102E). Equal aliquots of the lysates were supplemented with or without $Ca^{2+}$/calmodulin prior to immunoprecipitation.

D) Quantification of CaMKIIα and GFP-$Ca_V1.3$ signals in GFP-immune complexes from three independent transfection replicates. GFP-$Ca_V1.2$ signals were normalized to the input signal, while CaMKII signals were first normalized to GFP-$Ca_V1.2$ in the corresponding IP. Ratios within each replicate were normalized to the $Ca^{2+}$/calmodulin/CaMKIIα-WT condition to allow for pooling of the data. Statistical analysis: two-way ANOVA with Šídák's post hoc test.

Images of immunoblots for each protein in panels A and C were from the same transfection replicate gel/blot and were collected using identical detection sensitivity.

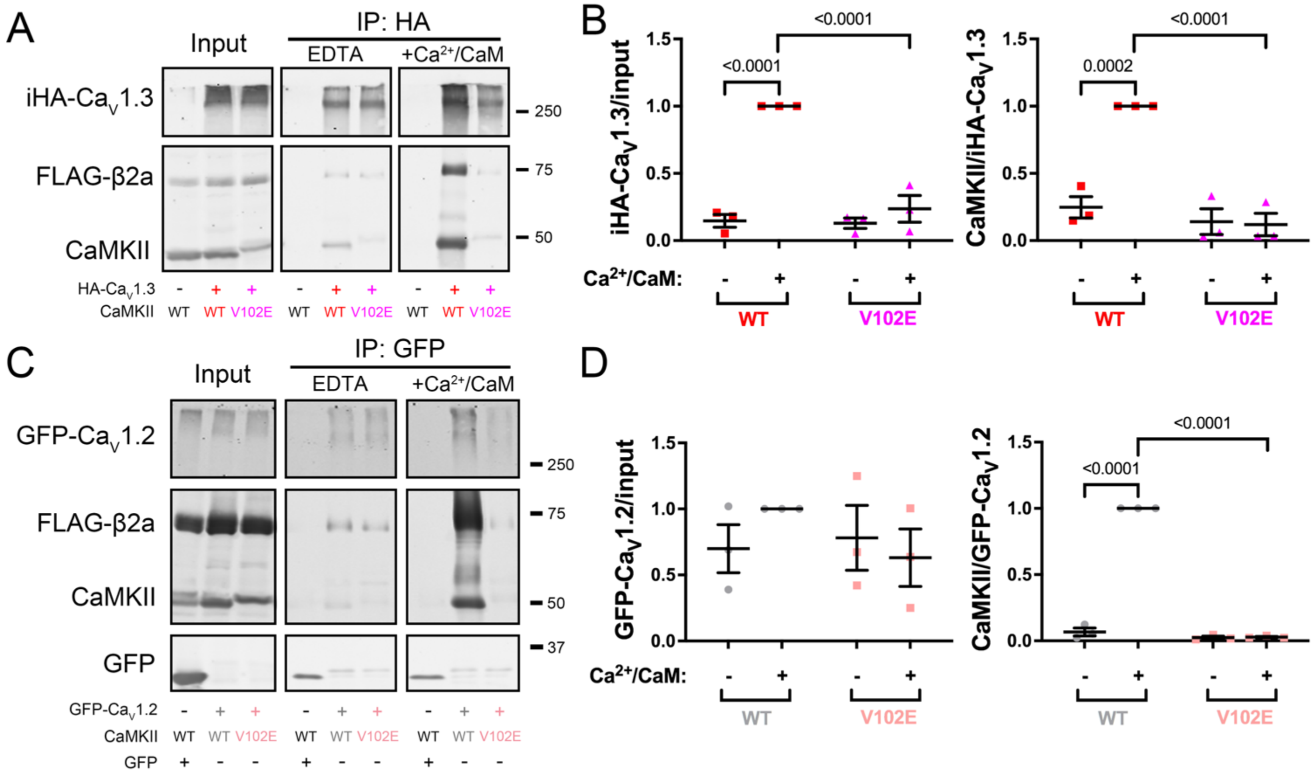

**Supplemental Figure 7 (related to Figure 4). Sensitivity of Shank3 interactions with other proteins to 2.5% 1,6-hexanediol.**

A) Representative immunoblots of the input and anti-GFP IPs from soluble fractions of HEK293T cells co-expressing GFP- and mApple-Shank3. The IPs were conducted in the absence or presence of 2.5% 1,6-hexanediol (1,6-HD).

B) Quantification of GFP and mApple signals in GFP-immune complexes from four independent transfection replicates. GFP-Shank3 signals in the IP lanes were normalized to the corresponding input (GFP-Shank3/input); mAplle-Shank3 signals were normalized to GFP-Shank3 in the corresponding lane (mApple-/GFP-Shank3). Ratios within each replicate were then normalized to the condition without 1,6-HD application within each replicate.

C) Representative immunoblots of the input and anti-HA IPs from soluble fractions of HEK293T cells co-expressing iHA-$Ca_V1.3$, mCherry-$Ca_V1.3$, α2δ, FLAG-β2a subunits, and GFP-Shank3. The IPs were conducted in the absence or presence of 2.5% 1,6-HD.

D) Quantification of iHA-$Ca_V1.3$, mCherry-$Ca_V1.3$, GFP-Shank3, and FLAG-β2a signals in HA-immune complexes from two independent replicates. Immunoblot signals were normalized as described in Figure 2, relative to the condition without HD application within each replicate.

E) Representative immunoblots of the input and anti-GFP IPs from soluble fractions of HEK293T cells co-expressing GFP-Shank3 and CaMKIIα after addition of $Ca^{2+}$/calmodulin. The IPs were conducted in the absence or presence of 2.5% 1,6-HD.

F) Quantification of GFP-Shank3 and CaMKIIα signals in GFP-immune complexes from four independent transfection replicates. Immunoblot signals were normalized similarly as described in panel B, relative to the condition without HD application within each replicate.

Statistical analyses: one sample t-tests.

Images of immunoblots for each protein in panels A, C, and E were from the same transfection replicate gel/blot and were collected using identical detection sensitivity.

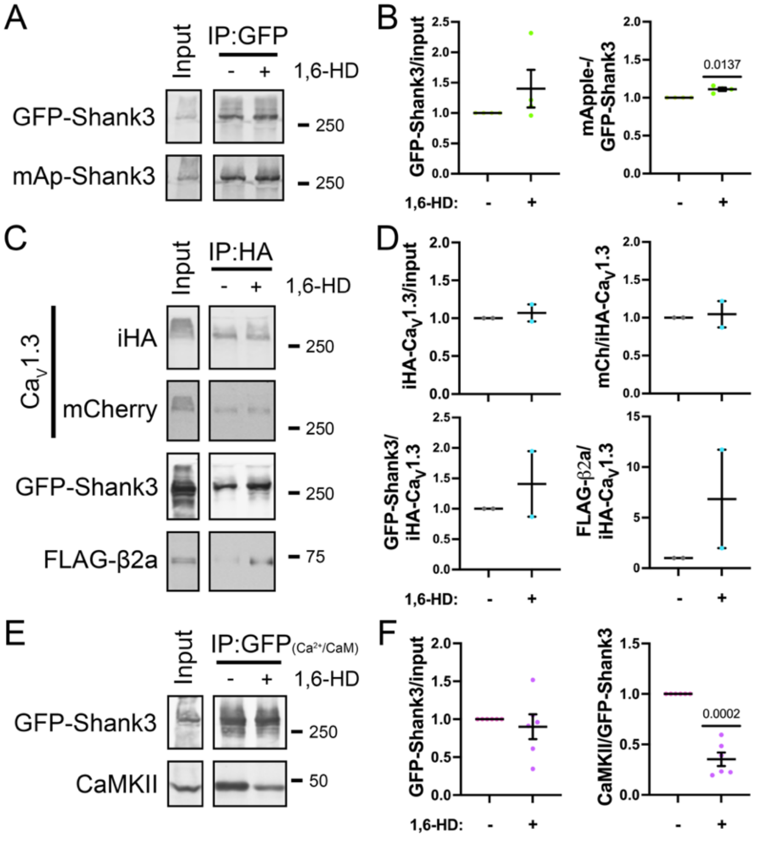

**Supplemental Figure 8 (related to Figures 6-7). Efficiency of CaMKII knockdown by CaMKII specific shRNAs.**
Primary rat hippocampal neuron cultures (14 DIV) were transfected to express $Ca_V1.3$ with an extracellular HA tag (sHA-$Ca_V1.3$), FLAG-β2a, and either GFP-nonsense shRNA (GFP-nssh) or GFP-CaMKIIα/β shRNA (GFP-CaMKII-sh). Neurons were fixed at DIV21, permeabilized and then immunostained for endogenous CaMKII.
A) Representative maximum intensity projection of the Z-stack of confocal GFP, DAPI and eCaMKII images merged using FIJI. The soma was outlined by the white dashed lines based on the GFP expression. Scale bar, 20 µm.
B) Quantification of somatic CaMKII staining intensity, normalized to CaMKII staining in nearby non-transfected (NT) cell somas on the same coverslips. Data from three independent transfections: total of 29 and 32 neurons for GFP-nssh, and GFP-CaMKII-sh, respectively.
B') Re-analysis of data in panel B by the 3 replicates.
Statistical analysis in B and B': unpaired t-test.

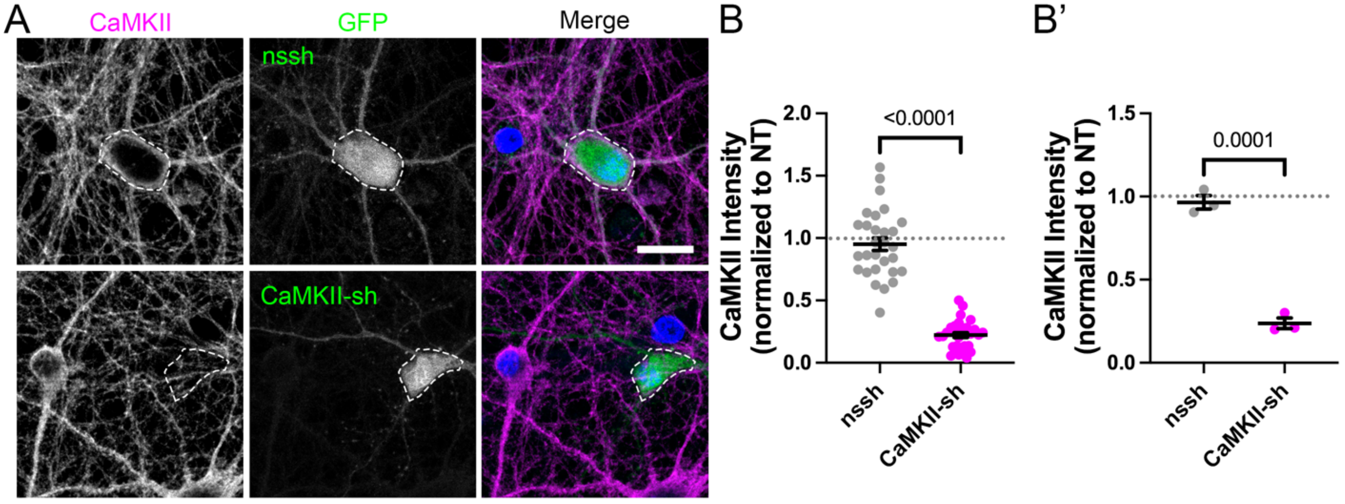

**Supplemental Figure 9 (related to Figures 6, 7). CaMKII is required for depolarization-induced CREB Ser133 phosphorylation in neurons over-expressing sHA-$Ca_V1.3$.**
Primary rat hippocampal neurons (DIV 14) expressing sHA-$Ca_V1.3$ and FLAG-β3 or -β2a with either GFP-nonsense shRNA (nssh) or GFP-CaMKIIα shRNA/CaMKIIβ-shRNA (CaMKII-sh) were incubated for 90 s with 40K/5K at DIV 21 and fixed (see Figure 1). Neurons were immunostained for the HA tag without permeabilization and then permeabilized for staining for DAPI and pCREB (see methods). Images were collected using confocal microscopy. Data for neurons expressing FLAG-β3 or -β2a were pooled because they were not significantly different.
A) Representative images of pCREB, DAPI, sHA, and GFP in neurons without (NT) or with transfection (nssh or CaMKII-sh). Scale bar, 5 µm.
B) Quantification of pCREB signal after 5K or 40K treatment in neurons with or without nssh expression (NT: n = 129 for 5K and 141 for 40K; nssh: n = 41 for 5K and 40 for 40K).
B') Re-analysis of data from panel B by averaging pCREB signal intensities in NT and nssh transfected neurons within each independent replicate (n=4 cultures/transfections).
C) Quantification of pCREB signal after 5K or 40K treatment in neurons with or without CaMKII-sh expression (NT: n = 113 for 5K and 204 for 40K; nssh: n = 41 for 5K and 50 for 40K).
C') Re-analysis of data from panel B by averaging pCREB signal intensities in NT and CaMKII-sh transfected neurons within each independent replicate (n=4 cultures/transfections, 2 each using FLAG-β3 and -β2a).
Statistical analyses: two-way ANOVA with with Šídák's post hoc tests.

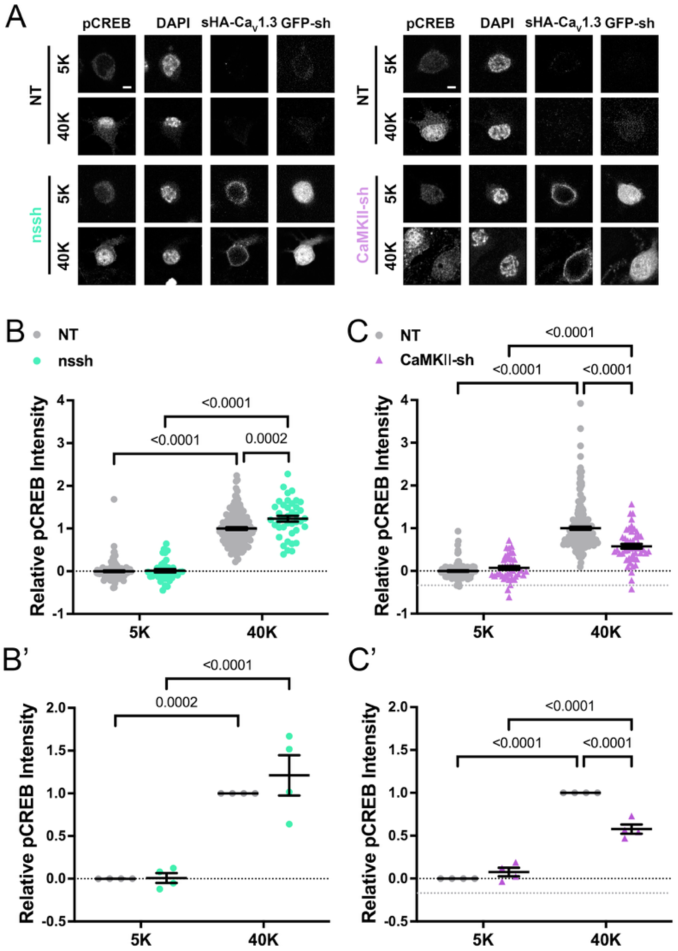

**Supplemental Figure 10 (related to Figure 6). Depolarization-induced $Ca_V$1.3 and $Ca_V$1.2 clustering in hippocampal neurons is disrupted by CaMKII knockdown.**

Hippocampal neurons expressing sHA-$Ca_V$1.3 or HA-$Ca_V$1.2 and FLAG-β2a with either GFP-nssh or GFP-CaMKII-sh (DIV 21) were incubated for 90 s with 40K/5K and fixed (see Figure 1). Neurons were immunostained for HA, permeabilized, and then immunostained for endogenous CaMKII (eCaMKII). Images collected using Airyscan super-resolution confocal microscopy.

A) Representative images of sHA-$Ca_V$1.3 and CaMKII staining in soma of neurons expressing GFP-nssh. Scale bar, 5 µm.

B-C) Quantification of sHA-$Ca_V$1.3 cluster intensity and density in GFP-nssh (B) and GFP-CaMKII-sh (C) expressing neurons (n = 41 (5K), 33 (40K) neurons expressing nssh; n = 34 (5K) and 28 (40K) neurons expressing CaMKII-sh). Data were quantified from 5 independent cultures/transfections and normalized to the mean for transfected neurons following the 5K incubation within each replicate. Statistical analyses: unpaired t-tests.

D) Representative images of sHA-$Ca_V$1.2 and CaMKII staining in soma of neurons expressing GFP-CaMKII-sh. Scale bar, 5 µm.

E-F) Quantification of sHA-$Ca_V$1.2 cluster intensity and density in GFP-nssh (E) and GFP-CaMKII-sh (F) expressing neurons (n = 24 (5K), 27 (40K) neurons expressing nssh; n = 27 (5K) and 24 (40K) neurons expressing CaMKII-sh). Data were quantified from 3 independent cultures/transfections and normalized to the mean for transfected neurons following the 5K incubation within each replicate. Statistical analyses: unpaired t-tests.

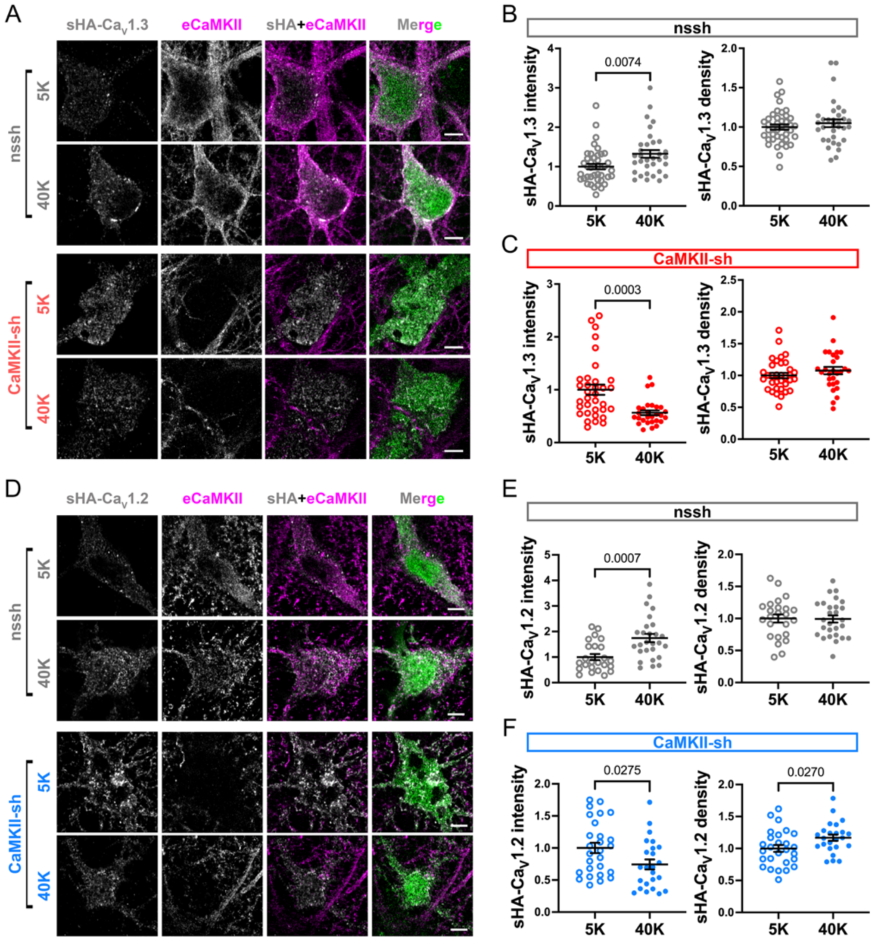

**Supplemental Figure 11 (related to Figure 6 and Supplemental Figure 9). Depolarization-induced Cav1.3 and Cav1.2 clustering in hippocampal neurons is disrupted by CaMKII knockdown.**

Re-analysis of data in Figures 6 and S9 by averaging sHA cluster intensity and density in soma (A and C) dendrite (B and D) of neurons co-expressing sHA-Cav1.3 (A and B) or sHA-Cav1.2 (C and D) and FLAG-β2a with nssh or CaMKII-sh within each independent replicate, then normalizing to values in 5K-treated nssh expressing neurons (n=5 cultures/transfections for sHA-CaV1.3 and 3 cultures/transfections for sHA-Cav1.2). Statistical analyses: one sample t-tests.

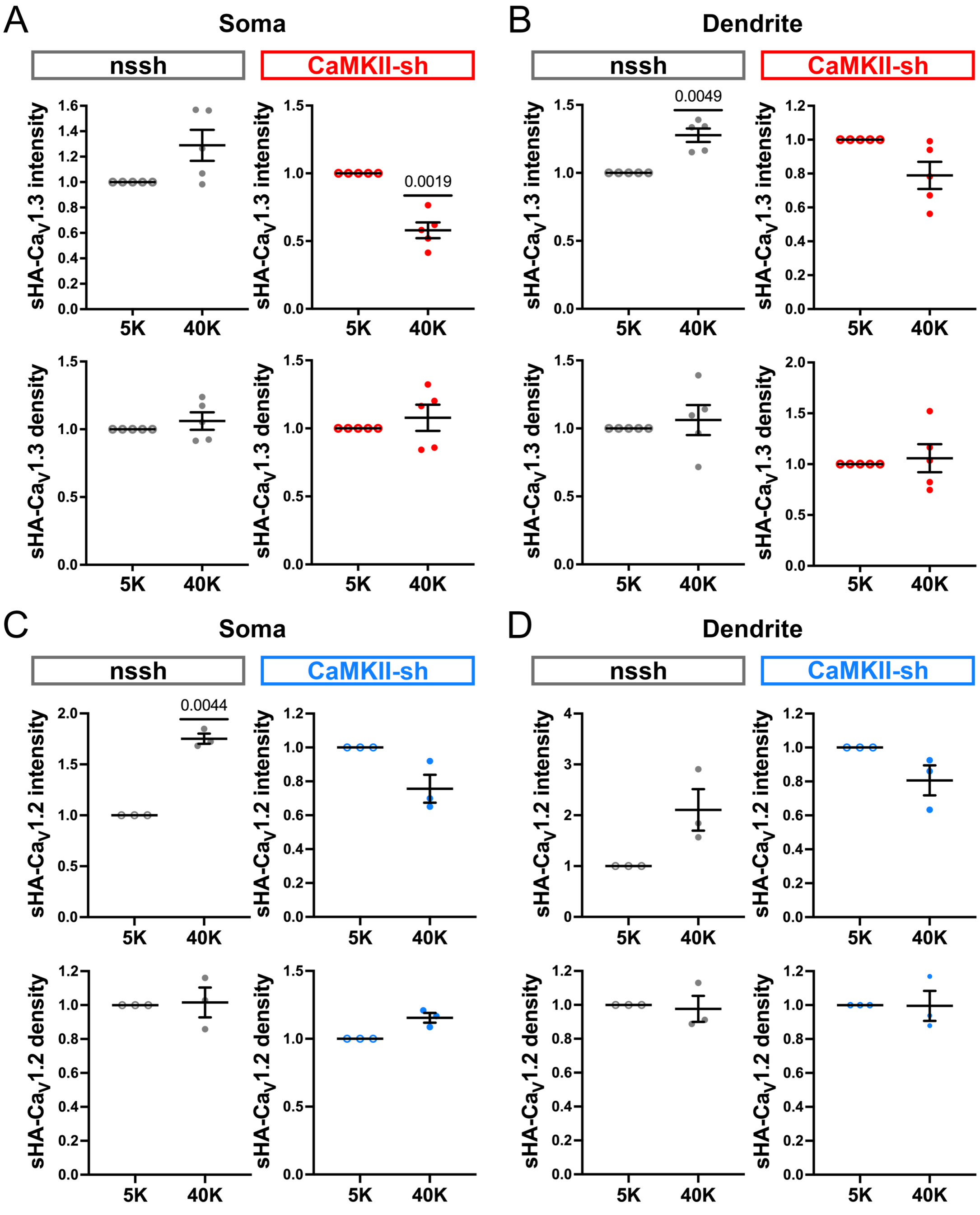

**Supplemental Figure 12 (related to Figure 7). Depolarization-induced $Ca_V1.3$-$Ca_V1.2$ co-clustering is CaMKII-dependent in hippocampal neurons.**
Hippocampal neurons expressing sHA-$Ca_V1.3$ and FLAG-β2a with either GFP-nssh or GFP-CaMKII-sh (DIV21) were incubated for 90 s with 40K/5K and fixed (see Figure 1). Neurons were immunostained for HA, permeabilized and then immunostained for endogenous $Ca_V1.2$ (e$Ca_V1.2$). Neurons were imaged using an Airyscan super-resolution confocal microscopy.
A) Representative images of sHA staining, e$Ca_V1.2$ staining, and merged images in soma of neurons expressing GFP-nssh or CaMKII-sh. Scale bar, 5 µm.
B) ROIs were defined based on sHA staining in order to measure the ratio of normalized e$Ca_V1.2$ intensity to normalized sHA intensity. Quantification of e$Ca_V1.2$ staining within sHA-$Ca_V1.3$ ROIs (see methods) in neurons from three independent transfected cultures. n = 20 (5K) and 21 (40K) neurons expressing nssh; n = 20 (5K) and 22 (40K) neurons expressing CaMKII-sh. Data were normalized to the average e$Ca_V1.2$/sHA-$Ca_V1.3$ ratio in nssh neurons with 5K treatment for each transfection.
B') Re-analysis of data in panel B by the 3 replicates.
Statistical analyses: two-way ANOVA with Šídák's post hoc tests.

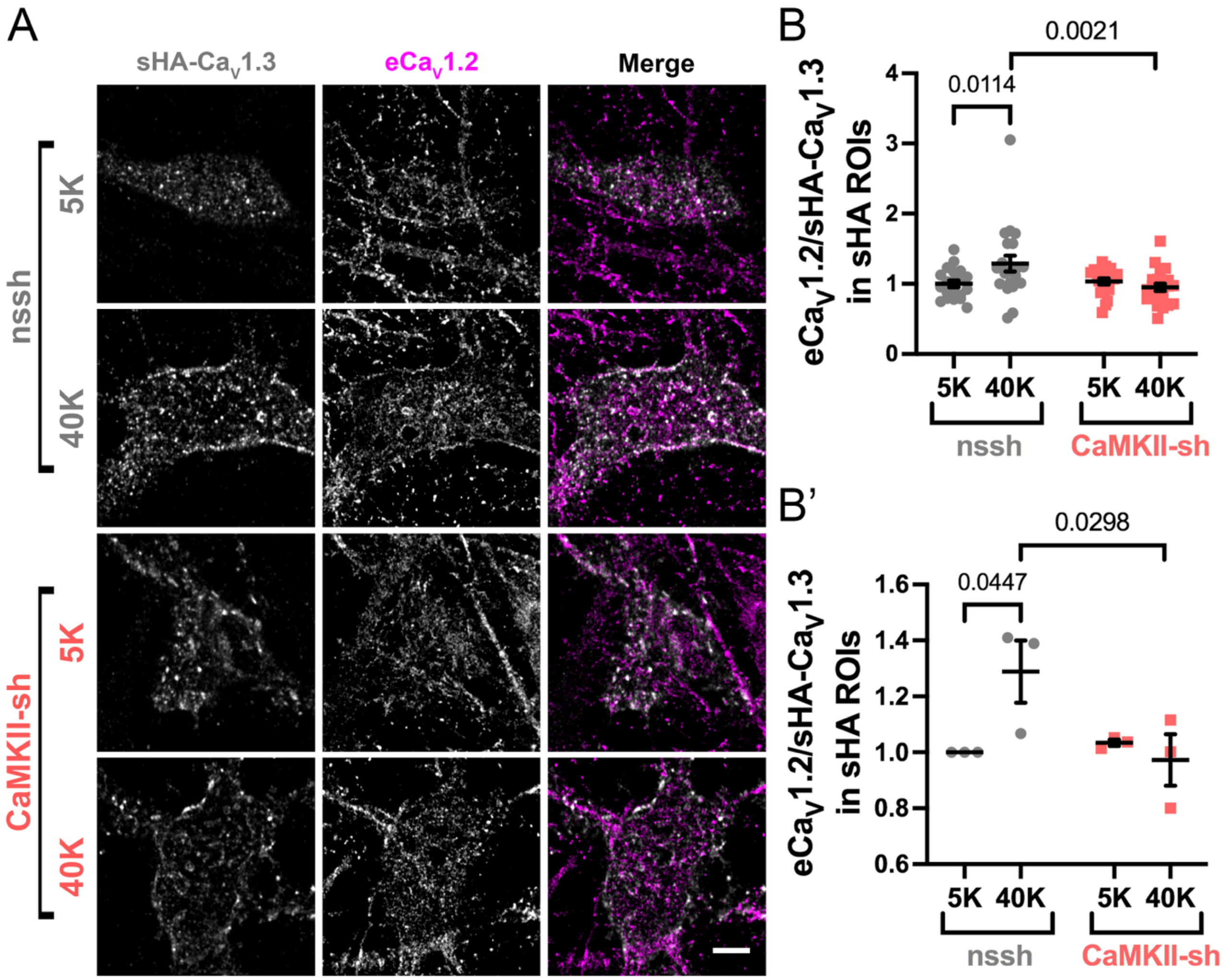
